# Lateral opening site of human oligopeptide transporter 2 plays a key role in the interaction with polymyxins

**DOI:** 10.64898/2026.04.01.715775

**Authors:** Xukai Jiang, Yining Luo, Mohammad A. K. Azad, Limei Xu, Min Xiao, Tony Velkov, Kade D. Roberts, Visanu Thamlikitkul, Qi Tony Zhou, Fanfan Zhou, Jian Li

**Affiliations:** National Glycoengineering Research Center, Shandong University, Qingdao 266237, China; Molecular Drug Development Group, Sydney Pharmacy School, Faculty of Medicine and Health, The University of Sydney, Sydney 2006, Australia; Biomedicine Discovery Institute, Infection Program, Monash University, Melbourne 3800, Australia; Division of Infectious Diseases and Tropical Medicine, Department of Medicine, Faculty of Medicine Siriraj Hospital, Mahidol University, Bangkok 10700, Thailand; Department of Industrial and Molecular Pharmaceutics, College of Pharmacy, Purdue University, West Lafayette, IN 47907, USA; Save Sight Institute, Faculty of Medicine and Health, The University of Sydney, Sydney 2006, Australia

**Author notes:** Correspondence: Professor Jian Li, Professor Fanfan Zhou.

**Keywords:** Antibiotic resistance, polymyxin-induced nephrotoxicity, hPepT2, structure-interaction relationship, drug discovery

## Abstract

Multidrug-resistant (MDR) Gram-negative bacteria pose a critical global health threat, while polymyxins remain a last-line therapy. However, their clinical use is limited by nephrotoxicity. Human oligopeptide transporter 2 (hPepT2) is a membrane transporter mediating the reabsorption of polymyxins in renal cells and contributes to their nephrotoxicity, but the molecular basis of their interaction remains unclear. Here, we investigated the structure-interaction relationship (SIR) of polymyxins with hPepT2 by integrating computational, chemical, and cell biology approaches. Bioinformatic modelling predicted an outward-facing hPepT2 structure and a potential transport pathway, with polymyxins interacting at the lateral opening, particularly E214, D215, D317, D342, and E622. Transporter mutagenesis and molecular analyses confirmed that D215 is critical for polymyxin binding, while other residues influence transporter turnover and/or expression. We subsequently synthesised polymyxin analogues with modifications at Dab1, Dab3, Dab5, and Dab9 of polymyxins, which reduced interactions with hPepT2. Notably, alanine substitution at Dab3 reduced nephrotoxicity in mice while retaining antibacterial activity. Overall, this proof-of-concept study demonstrates that the hPepT2-polymyxin SIR model provides a viable strategy for developing novel, safer lipopeptide antibiotics.

## Introduction

The World Health Organization has highlighted antimicrobial resistance as an urgent threat to global public health (1–3). In particular, the Gram-negative ‘superbugs’ *Acinetobacter baumannii, Pseudomonas aeruginosa*, and *Klebsiella pneumoniae* can develop resistance to nearly all available antibiotics, posing some of the most serious challenges in modern medicine (4). Due to the lack of effective alternative antibiotics, polymyxins are often used as a last-line therapy against these ‘superbugs’ (5). However, polymyxins can cause nephrotoxicity in up to 60% of patients after intravenous administration, which is the major dose-limiting factor limiting their clinical use (6, 7). Thus, there is an urgent need for safer, new-generation lipopeptide antibiotics (8).

It has been demonstrated that polymyxin-induced nephrotoxicity is associated with extensive reabsorption into renal proximal tubular cells (6). The intracellular concentration of polymyxins in renal proximal tubular cells can be 5,000-fold higher than the extracellular concentration, which consequently leads to oxidative stress, autophagy, cell cycle arrest, and apoptosis (6). Therefore, the extensive reabsorption of polymyxins into renal proximal tubular cells is the very first step in nephrotoxicity.

Human Solute Carrier transporters (SLCs) are the primary influx transporters responsible for the cellular uptake of many drugs, including antibiotics (9, 10). There are over 300 SLC isoforms, among which the SLCO, SLC22A, and SLC15A subfamilies are key in mediating drug transport across the cell membrane (11, 12). Our previous study revealed that the cellular uptake of polymyxins is mediated through hPepT2 in the kidney (13), which is encoded by a key isoform of the SLC15A subfamily, human oligopeptide transporter 2 *(SLC15A2)* (12). This transporter protein is highly expressed in renal tubular cells, mediating the uptake of polymyxins, with a comparable kinetic constant (*K*_m_, reflecting transporter-substrate binding affinity) to its prototype substrates (12–14). It is one of the proton-driven symporters, using the inwardly directed proton electrochemical gradient to drive the concentrative uptake of substrates across the cell membrane (15).

Substrates of hPepT2 include a range of di- or tri-peptides and peptide-like drugs, such as angiotensin-converting enzyme inhibitors (16). For instance, glycylsarcosine (Gly-Sar) is a synthetic dipeptide widely used as a model substrate for hPepT2, exhibiting a moderate binding affinity with a reported *K_m_* value of approximately 100 µM (17). Fluorescent peptide probes, such as 7-amino-4-methylcoumarin-3-acetic acid, have also been employed to evaluate hPepT2 function, with a reported *K_m_* value of approximately 780 µM (18). Several β-lactam antibiotics, including cefadroxil, have been identified as substrates of hPepT2, while antiviral agents such as valacyclovir can also be transported by this transporter (19, 20). Although many β-lactam antibiotics are substrates of hPepT2, some compounds, including cloxacillin and cefadroxil, have been reported to act as competitive inhibitors of hPepT2 (21). However, it remains unclear how hPepT2 recognises lipopeptide antibiotics, particularly polymyxins and this knowledge is critical for elucidating their interaction with polymyxins and its contribution to polymyxin-induced nephrotoxicity.

Here, we integrated computational and experimental approaches to elucidate the interaction between hPepT2 and polymyxins. Importantly, we conducted a proof-of-concept chemical biology study by designing novel lipopeptides and evaluating their cellular uptake, nephrotoxicity, and antibacterial activity. This study establishes the first structure-interaction relationship (SIR) model for polymyxins and hPepT2, laying the foundation for the rational drug design and discovery of novel lipopeptide antibiotics.

## Results

### MD prediction of the structural model of outward-facing hPepT2

We developed the three-dimensional structure of hPepT2 using AlphaFold2, as the outward structure of hPepT2 (Fig. 1a) has not been experimentally solved. The putative overall structure adopted an inward open conformation (Fig. 1b), which prevented the exploration of the initial capture of polymyxins from the extracellular side of the membrane. Parker *et al.* demonstrated the cryo-electron microscopy (cryo-EM) structure of rat PepT2 with an outward open conformation (22). As rat PepT2 shares 83% sequence similarity with hPepT2, it served as a suitable template for modelling the hPepT2 structure through homology modelling. Upon remodelling, the resultant structure of hPepT2 adopted an outward open conformation (Fig. 1c). Several transmembrane domains (TMs) are involved in the formation of the solvent-accessible cavity for polymyxin translocation, including TM1, TM2, TM5, TM7, TM8, and TM10. The gate residue pairs (L75-R329, P341-E79, D170-S635, and K642-A177) marking the conformations were clearly identified in the solvent-accessible regions on both the extracellular and intracellular surfaces (Fig. 1d). During the 500-ns all-atom MD simulations with the inward-facing conformation as an initial configuration, the distances between the extracellular gate residue pairs L75-R329 and P341-E79 increased from 20 Å to 31 Å, and from 15 Å to 21 Å, respectively. Meanwhile, the distances between the intracellular gate residue pairs D170-S635 and K642-A177 decreased from 15 Å to 13 Å, and from 13 Å to 10 Å, respectively (Fig. 1e). Notably, the transition from the inward conformation to the outward conformation uncovered by MD simulations is consistent with the homology model of rat PepT2 regarding the distance of the extracellular and intracellular gates (22). This observation indicates the accuracy of the proposed hPepT2 model.

**Figure 1.**
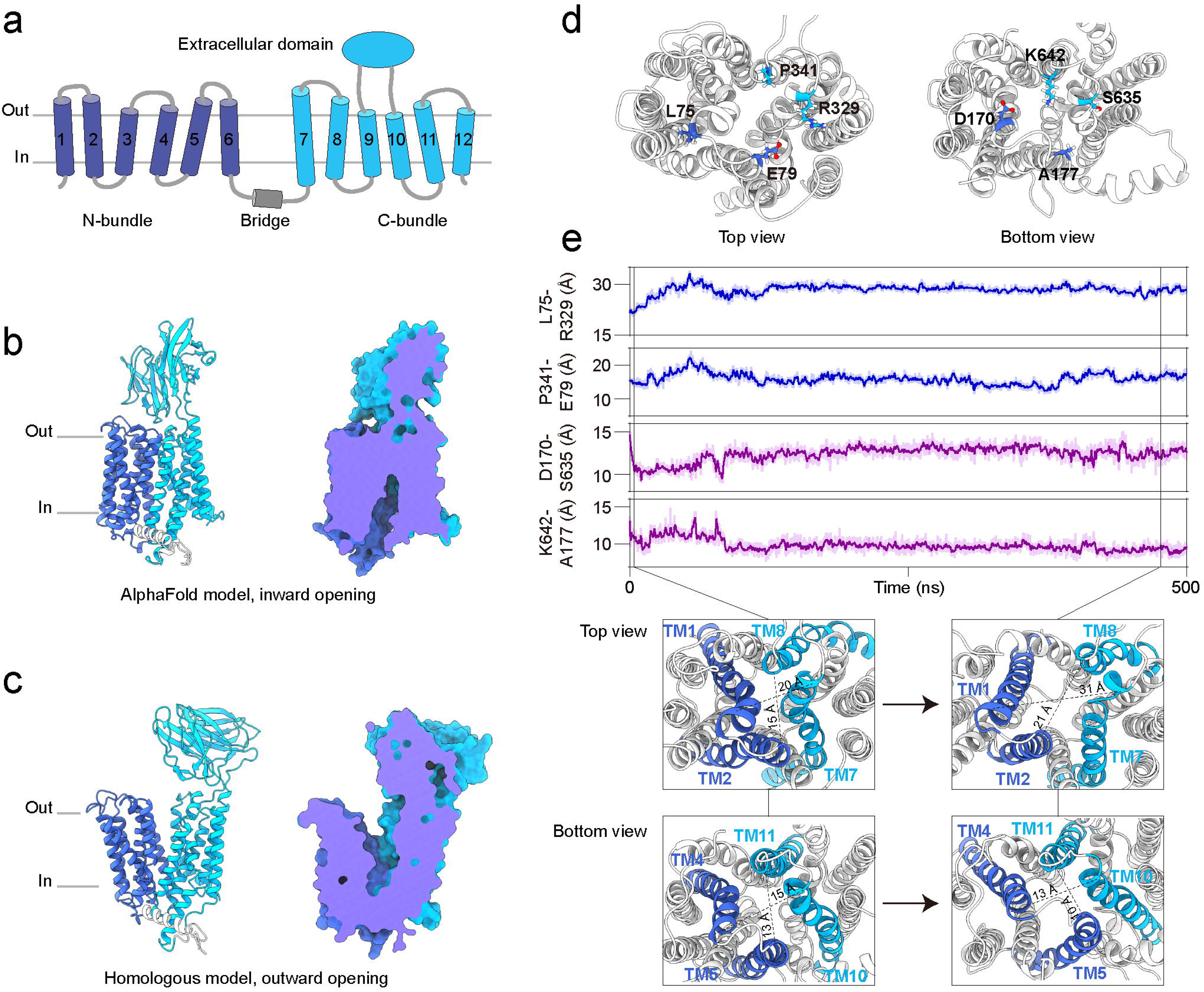
Structural conformations of hPepT2. **(a)** The structural topology of hPepT2. **(b)** The structural model of hPepT2 built with the AlphaFold approach. **(c)** The structural model of hPepT2 built with the homologous modeling method. The three-dimensional structure of rat PepT2 (PDB code: 7NQK) is used as the structural template. **(d)** The residues that are used to define the geometry of the channel of hPepT2. **(e)** The distances between the selected residues and representative structural snapshots are analyzed based on MD simulations.

### Polymyxin B binds to the lateral opening site of hPepT2 to enter the transport pathway

To further explore how hPepT2 recognises polymyxin molecules, we examined their interaction through long-timescale coarse-grained MD simulations. Although ten polymyxin B molecules were introduced into the simulation system, only one polymyxin molecule was observed to bind to the entrance of the hPepT2 binding pocket, while the other polymyxin molecules attached to the membrane lipids (Fig. 2a). Importantly, we discovered a distinct pathway by which polymyxins reach the extracellular lateral opening gate of hPepT2, as revealed by the simulation replicates (Fig. 2b and 2c), with distinct equilibrated binding conformations (Fig. 2c). Our MD simulations predicted a possible transport pathway for polymyxins via hPepT2. Initially, polymyxin B attached to the plateau of the TM1-TM6 helix bundle, mainly driven by electrostatic interactions between the Dab5, Dab8 and Dab9 residues of polymyxin B and the E79, D208, E214 and D215 residues of hPepT2 (Fig. 3). Subsequently, polymyxin B translocated to the junction between the TM7-TM12 helix bundle and the extracellular domains (ECDs) of hPepT2, where the Dab1, Dab3 and Dab5 residues of polymyxin B electrostatically interacted with the D342 and E555 residues of hPepT2. Polymyxin B then stabilised at the lateral opening gate of hPepT2 (i.e., lateral entry site), where the Dab8 and Dab9 moieties of polymyxin B electrostatically interacted with the D342 and E555 residues of hPepT2. Moreover, the hydrophobic atoms in Dab8 and Dab9 interacted with the F200, I201 and M204 residues of hPepT2, while the fatty acyl group of polymyxin B bound to the hydrocarbon tails of the phospholipid bilayer.

**Figure 2.**
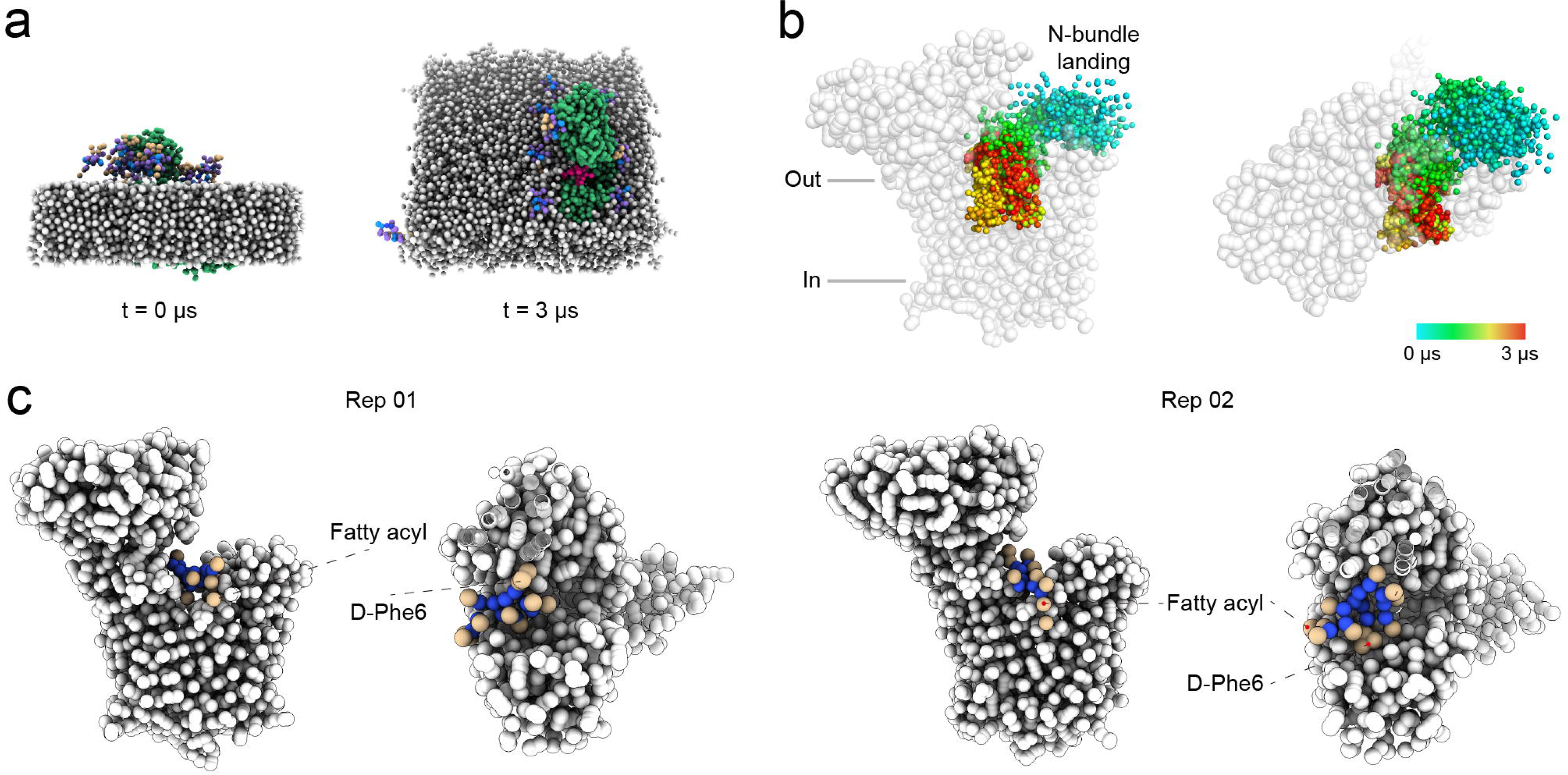
Spontaneous interactions of polymyxin B with hPepT2. **(a)** The interaction pathway of polymyxin B with hPepT2. The spatial locations of polymyxin B molecules are represented with colored spheres. The color spectrum from cyan to red indicates the simulation continuity from 0 to 3 μs. **(b)** The final state of two coarse-grained simulation replicates. hPepT2 is shown in grey spheres and polymyxin molecules are shown in brown and blue spheres. The fatty acyl group and D-Phe6 of polymyxin B are labelled. **(c)** The equilibrated conformation of the hPepT2-polymyxin complex according to the proposed pathway. The polymyxin B molecule is shown in brown and blue spheres with its D-Phe6, and *N*-terminal fatty acyl group labeled.

**Figure 3.**
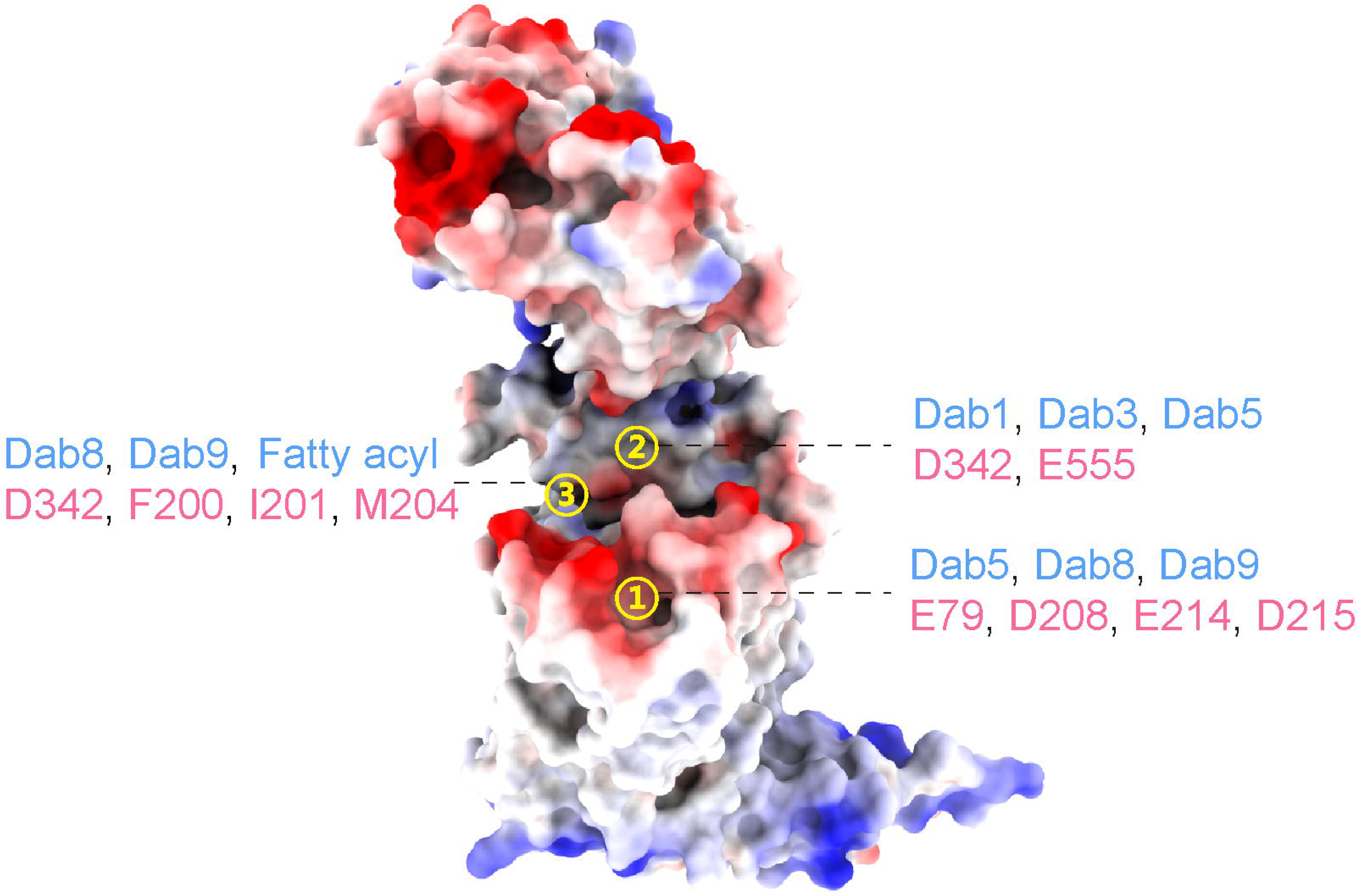
Key structural moieties and residues involved in the interaction of polymyxin B with hPepT2. The surface potential of hPepT2 is shown with red color indicating negative potential and blue color indicating positive potential.

To examine the detailed interactions between hPepT2 and polymyxin B, we performed all-atom MD simulations of the hPepT2-polymyxin B complex. During system preparation, the polymyxin B molecule was placed in four different poses within the complex to enhance the sampling of binding dynamics. The results revealed that Dab1, Dab8, and Dab9 form strong electrostatic interactions with hPepT2, with interaction energies ranging from -180 to -260 kJ/mol (Fig. 4), while Dab3 and Dab5 may also contribute significantly. Together with observations from the coarse-grained simulations, these results indicate that the positively charged Dab residues are key determinants of their binding affinity to hPepT2.

**Figure 4.**
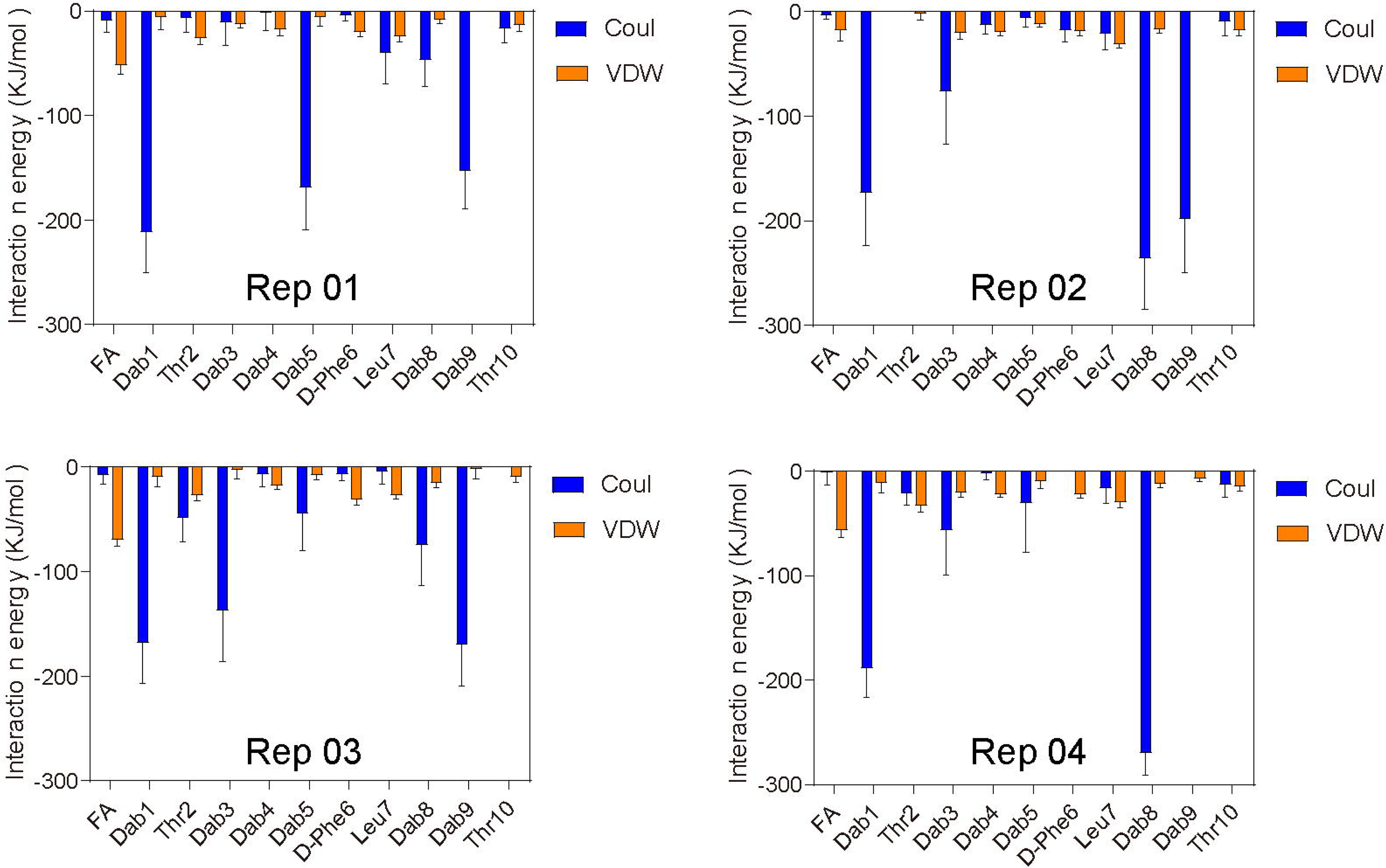
Interaction energy between hPepT2 and each residue of polymyxin B. The interaction energy was calculated using four MD simulation replicates. **Note:** Coul: electrostatic interaction; VDW: hydrophobic interaction.

### Functional mutagenesis elaborates on the predicted transport pathway

At physiological pH, there are five positively charged residues (i.e. Dab residues) on the polymyxin molecule, and we assumed that the key negatively charged residues of hPepT2 identified by the MD simulations (i.e. E79, D208, E214, D215, D317, D342, E555, and E622) may play crucial roles in the recognition of and interactions with polymyxins. We employed functional mutagenesis to confirm their roles in the uptake of polymyxins, as predicted by our MD studies. Through the commonly used alanine-scanning mutagenesis approach, we constructed the corresponding mutants of hPepT2. Notably, alanine substitution not only impacted the side-chain length of the proposed residues but also altered their charges from negative to neutral. It is postulated that such a drastic change in the residues of hPepT2 may substantially affect their interactions with polymyxins.

The transport activity of these hPepT2 mutants was evaluated by assessing the uptake of H^3^-Gly-Sar (a typical substrate of hPepT2) (Fig. 5a) (13) as well as the fluorescent polymyxin probe MIPS-9541 (Fig. 5b). The fluorescence images were also captured as a qualitative measurement of the overexpressing cells exposed to our dansyl-fluorescent polymyxin probe MIPS-9541, which indicated consistent results (Fig. 6). Our data showed that E79A, D80A, and D208A mutants maintained transport activity in the uptake of both substrates (Fig. 5). In contrast, E214A, D215A, D342A, and E622A mutants had impaired function in the transport of either substrate. The uptake of Gly-Sar was reduced to 65.4 ± 15.8%, 42.7 ± 13.2%, 49.4 ± 14.2% and 6.4 ± 4.7% of the wild type, respectively (Fig. 5a). In the case of MIPS-9541, the cellular uptake was decreased to 67.6 ± 14.2%, 38.9 ± 12.2%, 53.7 ± 16.2%, and 14.3 ± 8.0% of the wild type, respectively. Interestingly, D317A and E555A showed differential activity in the uptake of these substrates (33.1 ± 12.7% of the control for D317A and 93.6 ± 14.4% of the control for E555A in the uptake of Gly-Sar; 94.9 ± 13.3% of the control for D317A and 47.7 ± 16.2% of the control for E555A in the uptake of MIPS-9541), suggesting the distinct roles of both residues in interactions with different substrates.

**Figure 5.**
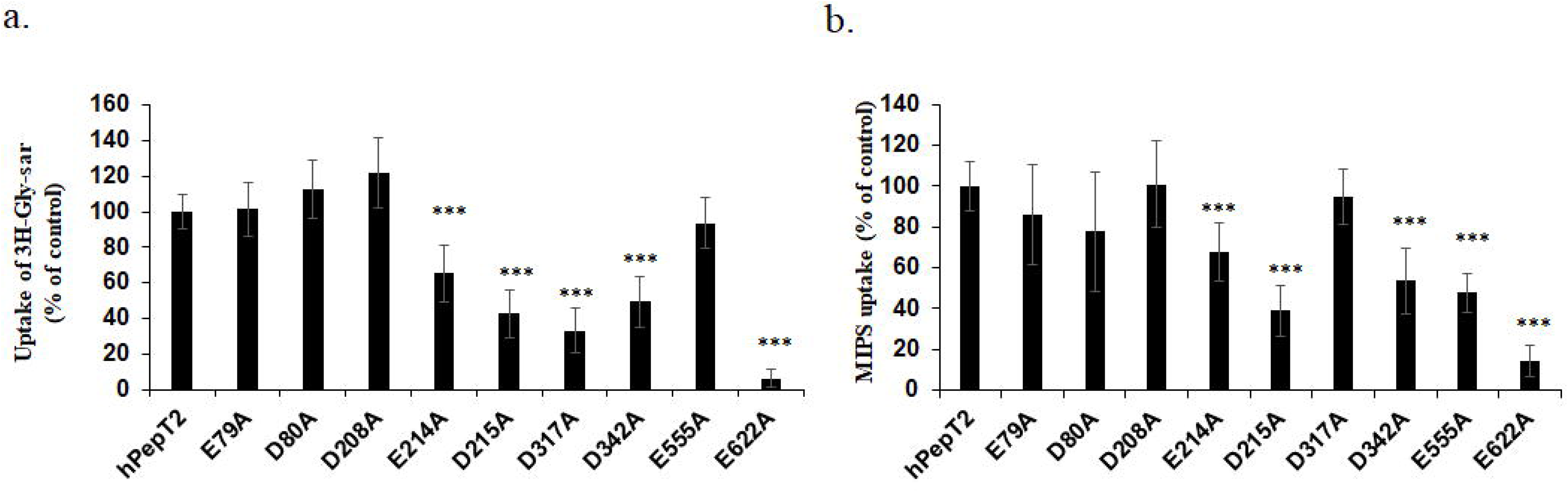
Uptake of ^3^H-Glycosarcosine and polymyxin B fluorescence probe MIPS-9541 by hPepT2 and its mutants. HEK293 cells were transfected with hPepT2 and its mutant constructs. **(a)** Uptake of [^3^H]-Gly-Sar (5 µM) by hPepT2 or its mutants. **(b)** Uptake of MIPS-9541 (10 µM) by hPepT2 or its mutants. Data are presented as mean ± SD. All experiments were performed in triplicate independently. \*\*\**p* < 0.001 *vs.* control by Welch’s *t*-test.

**Figure 6.**
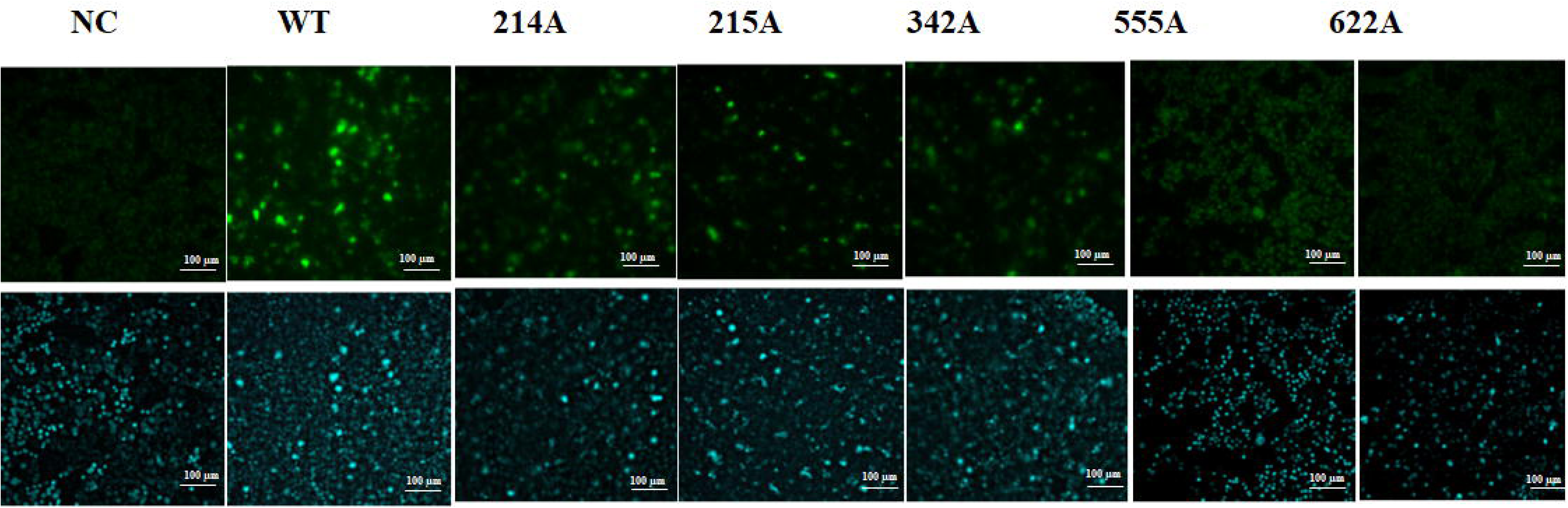
Uptake of the polymyxin B fluorescence probe MIPS-9541 in HEK293 cells overexpressing hPepT2 or its mutants assessed by fluorescence imaging. The fluorescence signaling of MIPS-9541 (green) was shown in the upper panel; the nuclei staining (blue) was shown in the bottom panel. Data are presented as mean ± SD. All experiments were performed in triplicate independently. Note: WT: wild type; NC: negative control.

Subsequently, kinetic analysis was conducted to characterise those hPepT2 mutants with impaired activity in the uptake of MIPS-9541 (i.e. E214A, D215A, D342A, and E555A). Notably, the E622A mutant was not included in this analysis due to its critically low transporter function. As shown in Table 1, the D215A mutant exhibited an approximately two-fold increase in both K_m_ (*p*=0.033, 95% confidence interval of 7.5 to 91.5 µM) and V_max_ (*p*=0.017, 95% confidence interval of 1510 to 7206 pmol/(µg/4 min)). It was also evident that the D342A mutant showed a mildly increased K_m_ (66.9 ± 13.2 µM of D342A *vs.* 47.1 ± 11.9 µM of the wild type, *p*=0.13, 95% confidence interval of -9.1 to 48.7 µM) without altering its V_max_. The V_max_ values of E214A and E555A were significantly reduced (2332 ± 24.2 pmol/(µg/4 min) of E214A, *p*=0.036, and 2376 ± 320.8 pmol/(µg/4 min) of E555A, *p*=0.022, *vs.* 4252 ± 652.9 pmol/(µg/4 min) of the wild type). However, the K_m_ values of E214A and E555A mutants were moderately decreased compared to those of the wild type (24.4 ± 4.5 µM of E214A, *p*=0.07 and 26.7 ± 7.0 µM of E555A, *p*=0.08 *vs.* 47.1 ± 11.9 µM of the wild type).

**Table 1.** Transporter kinetic parameters of MIPS-9541 uptake by hPepT2 and its mutants.

| Construct | K<sub>m</sub> (μM) | V<sub>max</sub> (pmol/(μg/4min)) |
| --- | --- | --- |
| hPepT2 | 47.1 ± 11.9 | 4252 ± 652.9 |
| E214A | 24.4 ± 4.48 | 2332 ± 24.2 * |
| **D215A** | **96.6 ± 20.9 \*** | **8610 ± 1363 \*** |
| D342A | 66.9 ± 13.2 | 4396 ± 583.1 |
| E555A | 26.7 ± 6.99 | 2376 ± 320.8 * |

\* $p < 0.05$ vs. control by Welch's *t*-test

Notably, a change in V_max_ value may be due to altered transporter protein expression and/or transporter turnover rate. Thus, the total cell and plasma membrane expression of hPepT2 and its mutants were investigated to further examine the molecular mechanism underpinning the functional alterations of these mutants. As shown in Fig. 7, E214A, D215A, E555A, and E622A significantly reduced total cell expression and cell surface expression. In the case of D342A, it retained transporter protein expression in the whole cell but had a markedly impaired cell surface expression.

**Figure 7.**
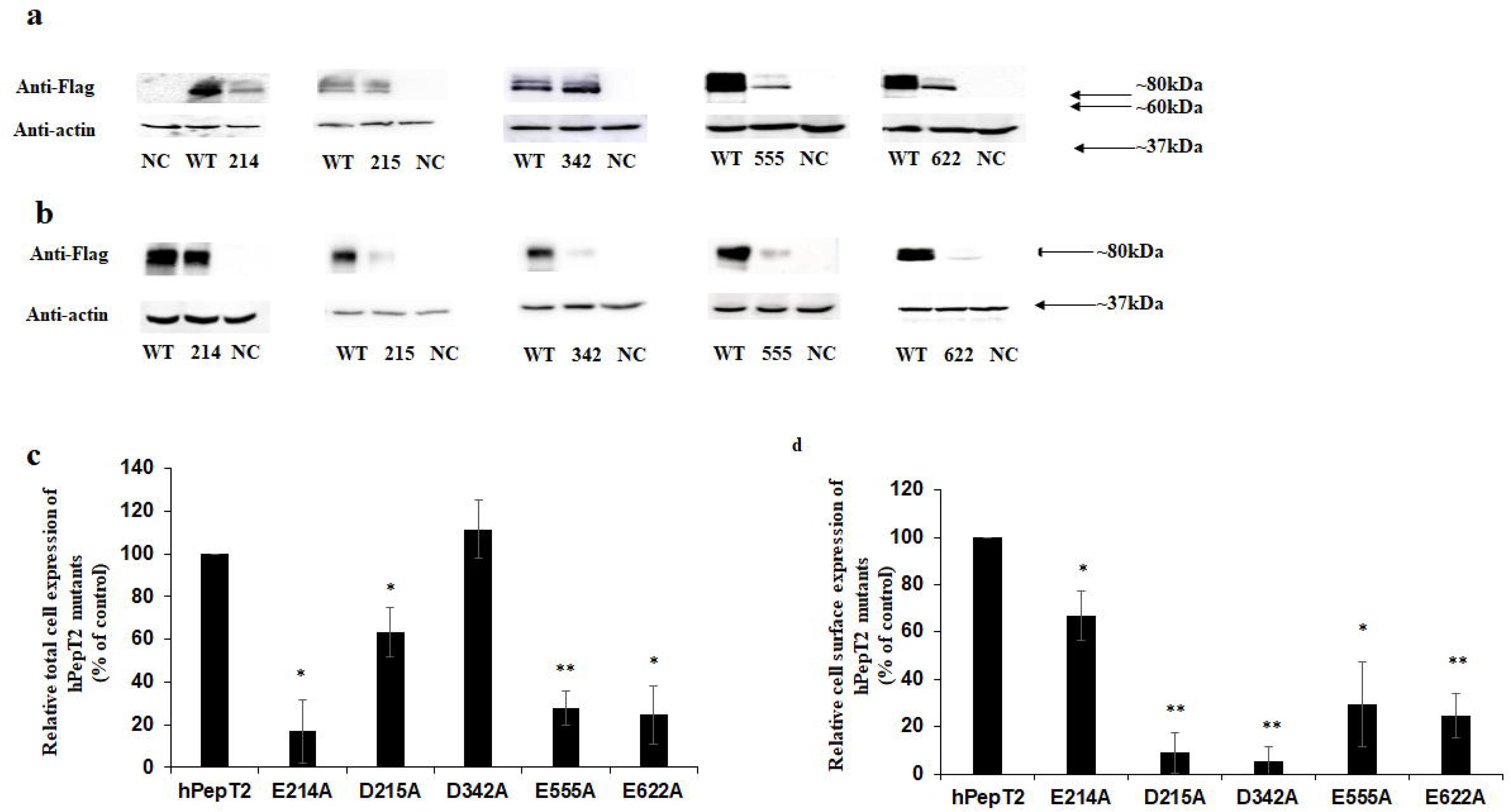
Total cell and cell surface expression of hPepT2-c-Flag and its mutants. HEK293 cells were transfected with hPepT2-c-Flag and its mutant constructs. **(a)** Cells were lysed and subjected to SDS-electrophoresis. Immunoblots were probed with anti-Flag antibody followed by probing with an anti-actin antibody as a loading control. Representative images of each hPepT2 mutant are shown. **(b)** Cell surface proteins were labelled with NHS-ss-Biotin and then pulled down with Streptavidin-agarose beads. Biotin-labelled surface protein samples were subjected to SDS-electrophoresis and the immunoblots were probed with anti-Flag antibody. Equal portions of the supernatant samples were separated by SDS-PAGE, and the immunoblots were incubated with anti-actin antibody. Representative images of each hPeT2 mutant are shown. **(c)** Densitometry analysis of the relative total cell expression of hPepT2 mutants (ratios of Flag/actin). **(d)** Densitometry analysis of the relative cell surface expression of hPepT2 mutants (ratios of Flag/actin). Data are presented as % of the hPepT2 wildtype control (mean ± SD). Experiments were repeated on three occasions. \**p* < 0.05; \*\**p* < 0.01 *vs.* wild-type control by unpaired *t*-test. Note: WT: wild type; NC: vector-transfected negative control.

### Chemical biology of polymyxin interaction with hPepT2

As described above, the Dab1, Dab3, Dab5, Dab8, and Dab9 residues of polymyxin B were predicted to be involved in the interaction with hPepT2. To test this hypothesis, we designed and synthesised five polymyxin B analogues with alanine substitutions on these Dab positions. We performed a transport uptake assay to evaluate their interactions with hPepT2. As shown in Fig. 8, the lipopeptide analogues FADDI-170, FADDI-175, FADDI-793, and FADDI-795 but not FADDI-167, showed reduced uptake in hPepT2-expressing cells, indicating that Dab1, Dab3, Dab5 and Dab9 but not Dab8, are likely involved in the interaction with hPepT2. Therefore, these four lipopeptide analogues may have less intracellular accumulation and reduced nephrotoxicity.

**Figure 8.**
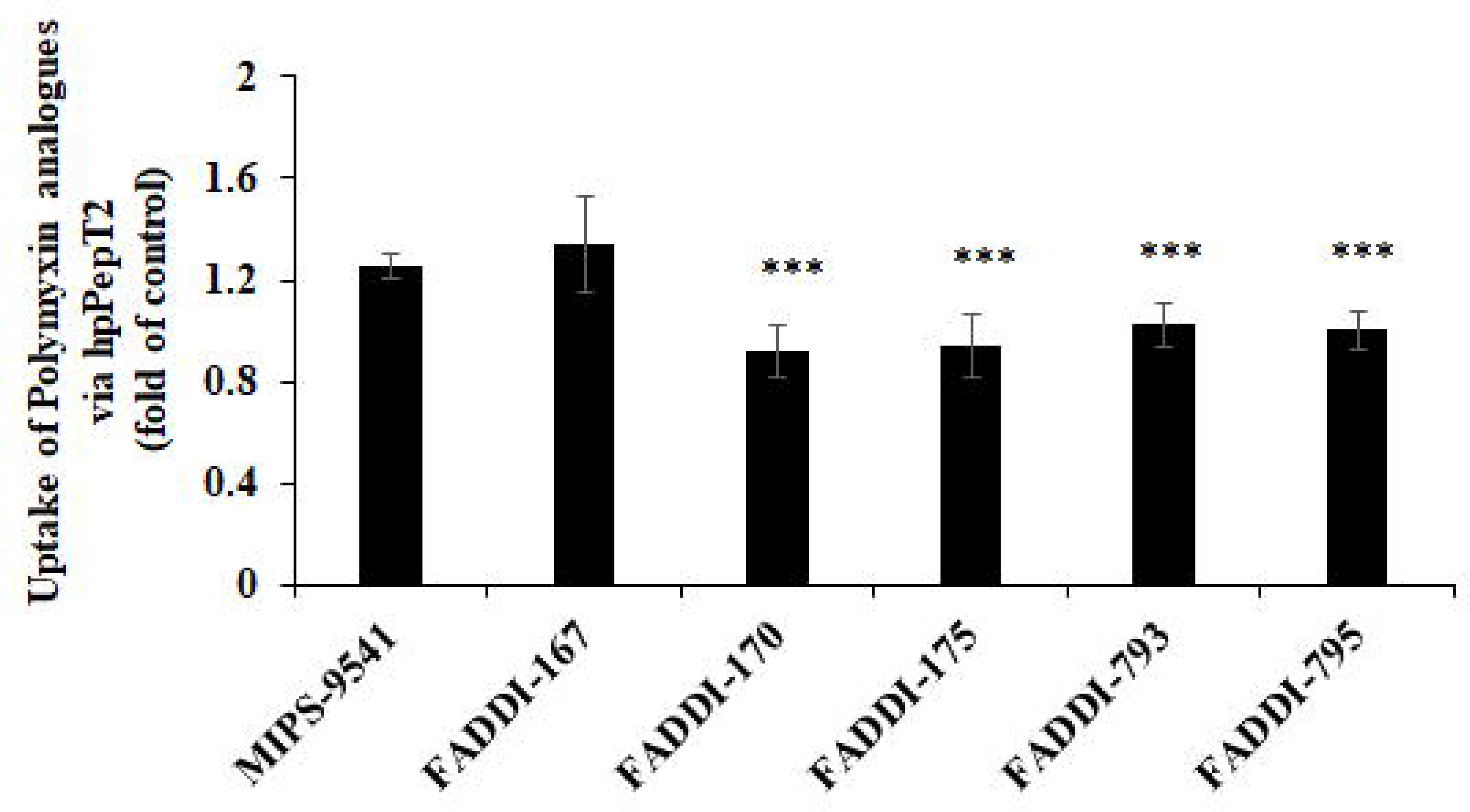
Uptake of the polymyxin B fluorescence probe MIPS-9541 and polymyxin analogues by hPepT2. The data are expressed as the fold change in the uptake via hPepT2 *vs.* that of the vector-transfected control. Data are presented as mean ± SD. All experiments were performed in triplicate independently. \*\*\**p* < 0.001 vs. MIPS-9541 control by Welch’s *t*-test.

### Pharmacological evaluations of lead polymyxin analogues

We firstly assessed the minimum inhibitory concentrations (MICs, the lowest concentration that inhibits visible growth of a microorganism after overnight culture) of the four lipopeptide analogues predicted to have disrupted interactions with hPepT2. As shown in Table 2, FADDI-170 showed significantly reduced activity against FADDI-PA025, FADDI-EC006, FADDI-EC003, FADDI-EC001, FADDI-AB034, as well as *A. baumannii* ATCC 19606 and ATCC 17978 (MICs increased ≥16-fold, compared to polymyxin B). FADDI-175 displayed low activity against all the tested strains (MICs ≥32 µg/mL), which was not further considered for subsequent testing. FADDI-793 was less active than polymyxin B against *A. baumannii* ATCC 19606, ATCC 17978, FADDI-AB034, FADDI-EC006, and FADDI-EC003. FADDI-795 was the most active among them, showing comparable MICs against these bacterial strains except for FADDI-PA025, when compared to polymyxin B. Interestingly, FADDI-793 and FADDI-795 were more active than polymyxin B against *K. pneumoniae* ATCC 13883. We then employed the mouse model established in our laboratory (23) to examine the nephrotoxicity of the four antibacterial polymyxin analogues. Interestingly, FADDI-795 showed no observable nephrotoxicity, while the other three analogues possessed mild kidney toxicity, comparable to that of polymyxin B (Table 2).

**Table 2.**
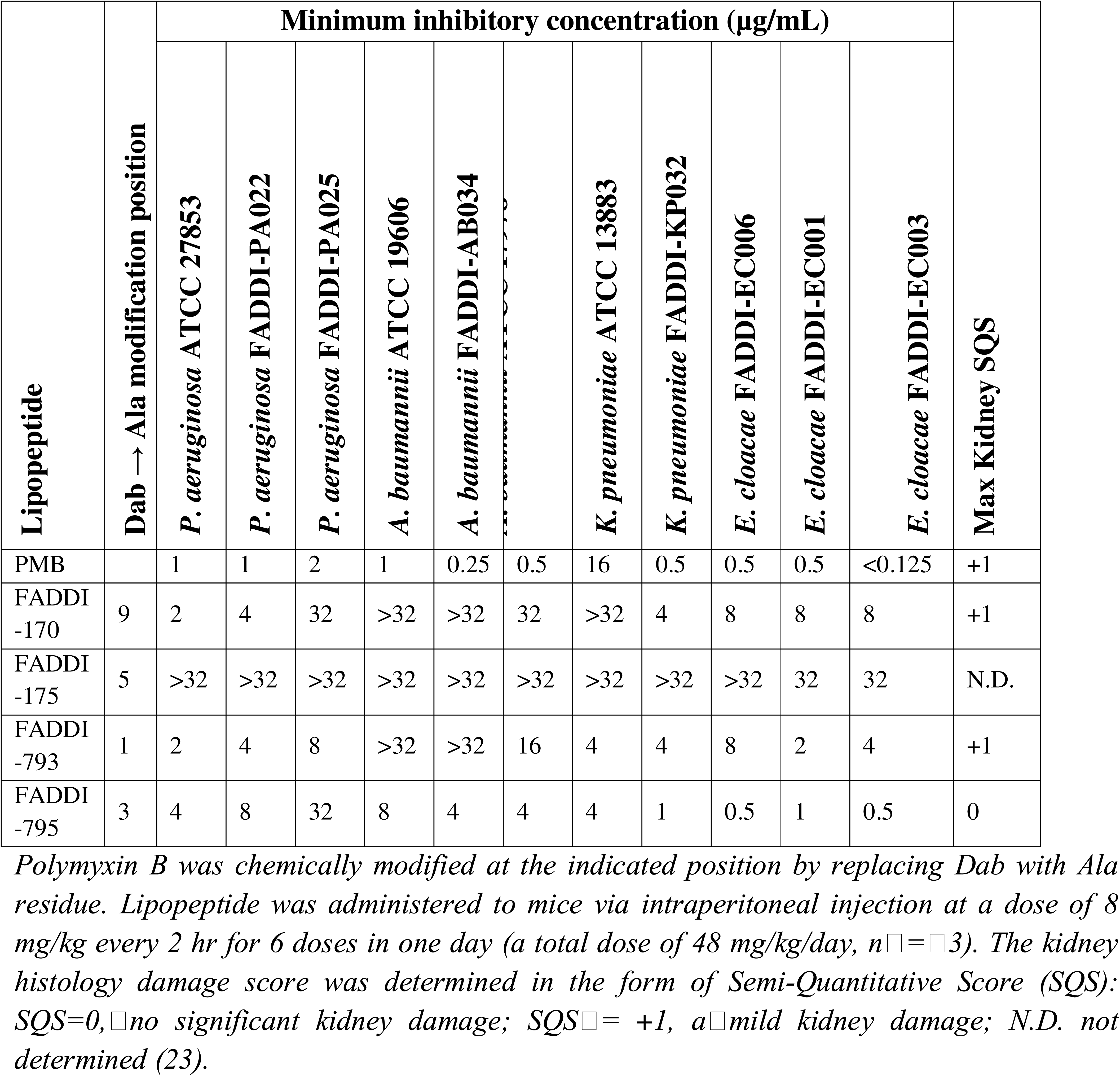
Chemical modifications, minimum inhibitory concentrations and nephrotoxicity of polymyxin B (PMB) and its analogues.

Overall, the transporter uptake study and pharmacological assessments of these polymyxin analogues provide critical mechanistic data on the interaction and transport of polymyxins by hPepT2. Importantly, our integrated computational and cell biology approach highlighted the feasibility of reducing polymyxin nephrotoxicity by disrupting their interactions with hPepT2. Moreover, hPepT2 is a viable target for designing novel polymyxin-like lipopeptide antibiotics with reduced nephrotoxicity.

## Discussion

Life-threatening MDR bacterial infections rank among the world’s most pressing health threats (24), yet the discovery of antibiotics with novel mechanisms of action has stalled for decades. Polymyxins remain a last-line therapy for MDR Gram-negative bacterial pathogens, particularly in low- to middle-income countries; however, their clinical use has been significantly limited due to nephrotoxicity (5, 6).

Our previous study demonstrated that polymyxins are substrates of hPepT2, which mediates the substantial reabsorption of polymyxins in renal proximal tubular cells, leading to nephrotoxicity (13). However, little is known about the interaction between polymyxin molecules and hPepT2. Notably, previous studies have reported that megalin mediates the cellular transport of polymyxins in the kidney via a receptor-mediated endocytosis mechanism (6, 25). Receptor-mediated endocytosis is closely linked to changes in signalling pathways; thus, disrupting the interaction between polymyxins and megalin may lead to potential signalling consequences and complicate treatment outcomes. In contrast, transporter-mediated uptake does not involve such signalling changes, making manipulation of polymyxin-hPepT2 interactions a more favourable strategy for mitigating polymyxin-associated nephrotoxicity. Furthermore, our previous study demonstrated a direct structural interaction of polymyxins with the membrane of human kidney proximal tubular cells (26), showing that polymyxin B inserts its fatty acyl tail into the hydrophobic region of the cell membrane. The stereochemistry at position 3 and the hydrophobicity at positions 6 and 7 of polymyxin B were found to be critical for this interaction. These findings provide important insights into how polymyxins alter membrane dynamics and lipid organization in kidney cells, contributing to a better understanding of polymyxin-induced nephrotoxicity.

The current study developed the first SIR model of polymyxins with hPepT2, expediting the discovery of novel safer lipopeptide antibiotics. We firstly employed MD simulations to predict the binding of polymyxins to hPepT2 (Fig. 1 and 2) and identified several critical amino acid residues of hPepT2 in the recognition of polymyxins. An outward-facing model of hPepT2 was first constructed, indicating the access of polymyxin molecules to the lateral opening gate of hPepT2 (Fig. 3). Limited by the sampling capability of MD simulations, the complete translocation pathway of polymyxin B through hPepT2 was not observed. The distinct conformation of polymyxin B indicates that it is situated in the central binding cavity of hPepT2. To further explore the conformational change of polymyxin molecules required for hPepT2 transport, all-atom MD simulations were conducted by varying the orientations of polymyxin B. Our results revealed that this conformational transition involved a notable flip of polymyxin B, which enhanced the interaction between its fatty acyl chain and the deeper regions of the predicted hPepT2 binding pocket. Notably, the subsequent transporter mutagenesis and molecular characterisations of hPepT2 explored our proposed SIR model. Our integrated computational and experimental findings provide a proof-of-concept that disrupting the interaction between polymyxin and hPepT2 is a novel strategy to reduce their nephrotoxicity.

As polymyxins are cationic at physiological pH, we strategically selected the negatively charged amino acids of hPepT2 (i.e. aspartic acid and glutamic acid) for further investigation. The significantly reduced uptake of polymyxin B by hPepT2 mutants E214A, D215A, D342A, E555A, and E622A demonstrated the critical role of these negatively charged residues in transporting polymyxins (Fig. 5 and 6). Moreover, the kinetic results of these transporter mutants elucidated the mechanism underpinning such functional impairment (Table 1). As both transporter cell surface expression and turnover rate account for the maximum velocity V_max_ of a transporter protein (12), it was critical to evaluate the protein expression of these transporter mutants in total cell lysates and at the cell surface (Fig. 7). Both the kinetic analysis and protein expression profiles (Table 3) showed different mechanisms of impaired polymyxin uptake by hPepT2. For example, it is highly likely that E214A reduced V_max_ due to reduced total cell and membrane protein expression. D215A reduced polymyxin uptake mainly due to impaired binding affinity to polymyxins (i.e. increased K_m_). The transporter turnover rate of D215A might increase to compensate for the reduced cell surface expression; thus, its V_max_ was overall increased. Similarly, the decreased polymyxin uptake mediated via D342A may be primarily attributed to its reduced polymyxin binding affinity. Its decreased membrane protein expression and likely elevated transporter turnover rate may have resulted in an unchanged V_max_. In contrast, the dysfunction of E555A may be mainly due to its reduced V_max_, likely resulting from a reduced total cell and membrane protein expression, although its binding affinity to polymyxins is slightly increased (i.e. decreased K_m_). The kinetic parameters of E622A were unable to be determined due to its low polymyxin uptake. Our data also indicated that its low protein expression in total cell and at cell surface may contribute to its impaired transport activity in polymyxin uptake. Notably, reduced total cellular expression of membrane transporters may result from impaired protein stability (27). In addition, cell surface transporter proteins undergo constitutive subcellular trafficking processes, including internalisation, recycling, and membrane targeting (28). Therefore, decreased cell surface expression of transporter proteins may arise from reduced total cellular expression and/or disrupted subcellular trafficking (27). Our findings suggest that E214, D215, E555, and E622 residues may play important roles in maintaining hPepT2 protein stability, whereas substitution of these residues with alanine may influence the subcellular trafficking of hPepT2. Nevertheless, the detailed molecular mechanisms underlying the changes in hPepT2 protein expression associated with these residues warrant further investigation but are beyond the scope of the current study.

**Table 3.** The kinetic parameters and protein expression profiles of hPepT2 mutant constructs.

| hPepT2 mutant | $K_m$ | $V_{max}$ | Total cell expression | Cell surface expression |
| --- | --- | --- | --- | --- |
| E214A | ↓ | ↓↓ | ↓↓ | ↓↓ |
| D215A | ↑↑ | ↑↑ | ↓↓ | ↓↓ |
| D342A | ↑ | *unchanged* | *unchanged* | ↓↓ |
| E555A | ↓ | ↓↓ | ↓↓ | ↓↓ |
| E622A | N.D. | N.D. | ↓↓ | ↓↓ |

↓: mildly or moderately decreased; ↓↓: significantly decreased; ↑: mildly or moderately increased; ↑↑: significantly increased; N.D.: not determined.

Notably, because some transporter mutants (e.g., D342A) exhibited both increased K_m_and reduced membrane expression, it is not possible to conclusively determine whether the decreased transporter function results from reduced binding affinity, lower protein expression, or a combination of both. This is a limitation of the current study. Nevertheless, our molecular assay results support the applicability of the bioinformatic predictions, as they allow us to focus the analysis on a limited set of transporter residues most relevant to our SIR model.

To validate the MD predictions, we further investigated the transport function of these mutant transporters through their uptake of the classic hPepT2 substrate Gly-Sar (Fig. 5a). The uptake of Gly-Sar was dramatically decreased in the mutants E214A, D215A, D317A, D342A, and E622A, which is consistent with the reduced cell surface expression of E214A, D215A, D342A, and E622A (Fig. 7). D317A is exceptional as it showed decreased uptake of Gly-Sar but maintained transport function in the case of polymyxin B, which indicated that the residue D317 may be only important for the uptake of Gly-Sar but not polymyxins. E555A is another exception, which remained functional for the uptake of Gly-Sar but reduced uptake of polymyxins. As E555A had a reduced total cell and cell surface expression, its increased apparent affinity for Gly-Sar may partially compensate for the reduced V_max_. Thus, the binding determinants of polymyxins may not fully overlap with those of Gly-Sar. Therefore, the transport inhibition study was only employed as a positive control initially and not considered in the subsequent evaluation of the antibacterial activity of polymyxin analogues.

According to the predicted topology model of hPepT2 (12), E214A and D215A are situated at the ECD between TM5 and TM6. D342A resides at the ECD between TM7 and TM8. E555A is located at the ECD between TM9 and TM10. E622A is situated within TM10. It was reported that the large ECD between TM9 and TM10 interacts with TM1 to maintain transport function (29). In our study, we demonstrated that substitution of E555 mildly increased the apparent binding affinity for polymyxin B while reducing V_max_, likely resulting from decreased protein expression. These findings highlight the need for more in-depth mechanistic understanding of hPepT2-mediated peptide transport. Notably, E622A in TM10 has a very low transport function in the uptake of polymyxin B, which is largely due to its low protein expression in the total cell and on the cell surface (Fig. 5 and 7). Therefore, this residue appears to be essential for preserving hPepT2 stability. Similar findings have been reported in other SLCs (27, 30, 31). According to the protein sequence alignment between hPepT2 and another key isoform of the SLC15A subfamily, hPepT1, E595 of hPepT1 is the equivalent residue to E622 in hPepT2. It has been shown that the alteration of E595 residue to cysteine substantially impacted transport function but not protein expression (32). This finding reflects the differing substrate specificities and transport capacities of hPepT1 and hPepT2 (12), highlighting the essential role of hPepT2 in mediating polymyxin transport.

Upon validating the predictions of the SIR model of polymyxins and hPepT2, our study was advanced to employ a chemical biology approach to rationally design and in-house synthesize polymyxin analogues. With the hope of disrupting the interaction between hPepT2 and these analogues, we aimed to discover candidate lipopeptide antibiotics with reduced nephrotoxicity while maintaining similar or better antibacterial activity. Based on the SIR model, the Dab1, Dab3, Dab5, Dab8, and Dab9 residues of polymyxins are likely to be involved in their interaction with hPepT2 (Fig. 4). Thus, we synthesised FADDI-167, FADDI-170, FADDI-175, FADDI-793, and FADDI-795 with alanine substitutions at these Dab residues. We first examined the hPepT2-mediated uptake of these polymyxin analogues. Interestingly, we discovered that all analogues except FADDI-167 disrupted the uptake via hPepT2 (Fig. 8). This finding indicated that FADDI-170, FADDI-175, FADDI-793 and FADDI-795 possibly impaired their uptake via hPepT2, which may contribute to reduced renal accumulation and nephrotoxicity. Thus, these four analogues were subjected to further pharmacological evaluations. Interestingly, FADDI-795 and FADDI-175 showed the best and least antibacterial activity among them, respectively (Table 4). Finally, we employed our mouse model to evaluate the nephrotoxicity of these lipopeptides. Using histological evaluations, FADDI-795 showed no observable renal toxicity in this mouse model and emerged as the lead compound with comparable antibacterial activity. A limitation of this research is that the nephrotoxicity assessment of new lipopeptides was performed in only a small number of mice in our proof-of-concept study. Nevertheless, the findings indicate that FADDI-795 is a promising candidate that will undergo further pharmacological evaluations to explore its translational potential, such as comprehensive pharmacokinetic studies, renal accumulation assessment and efficacy tests in relevant infectious disease models.

**Table 4.**
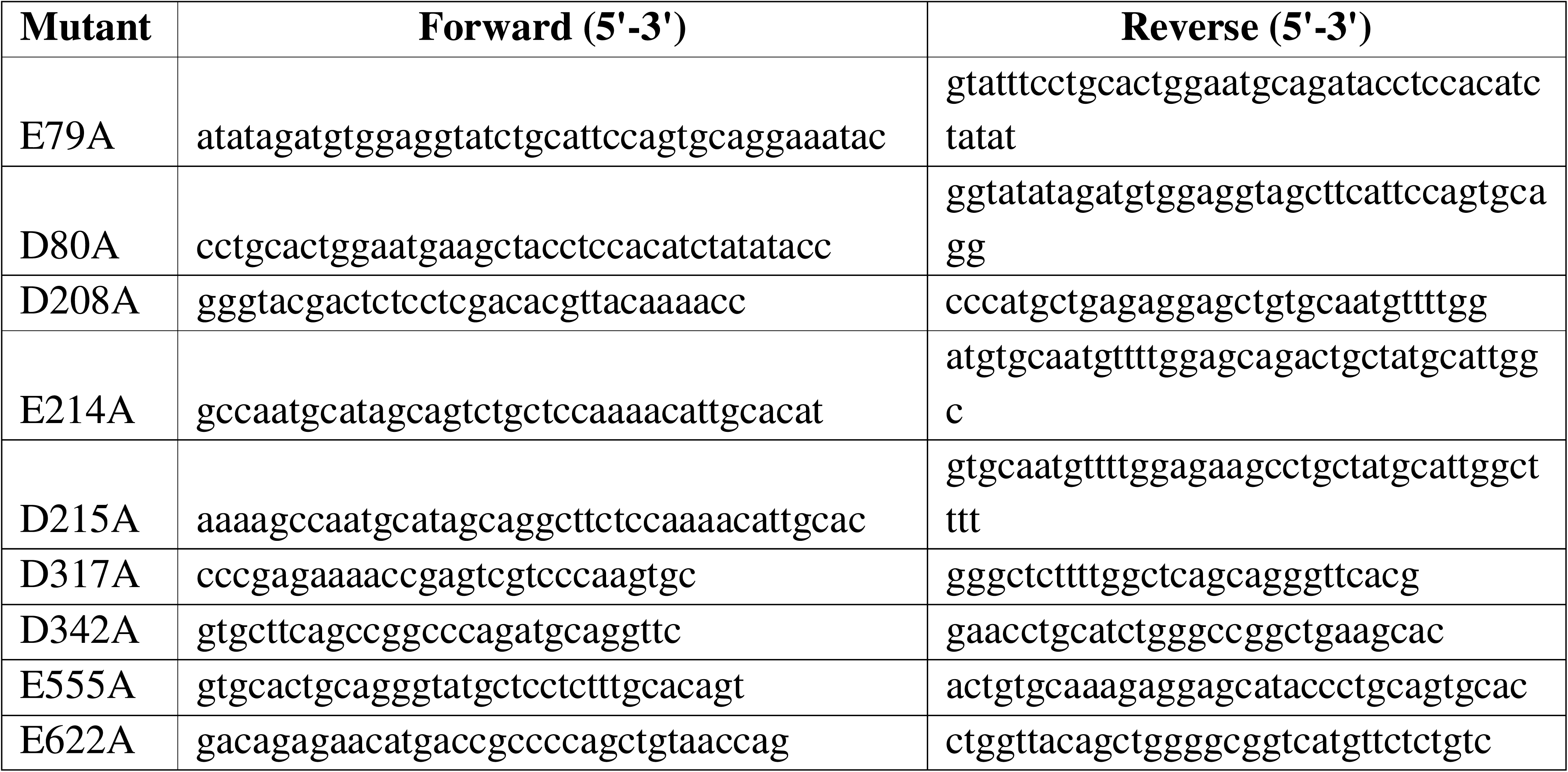
Primers used for hPepT2 mutagenesis.

Overall, our study developed the first SIR model of polymyxins with hPepT2 by integrating computational, chemical biology, and molecular approaches. Our molecular studies provide the first experimental validation of the predicted polymyxin-hPepT2 binding model. These mechanistic insights are essential for elucidating the mechanism of dose-limiting nephrotoxicity associated with polymyxin treatment. Moreover, our ongoing drug discovery efforts will further refine this SIR model and expedite the discovery and development of novel, safer lipopeptide antibiotics against Gram-negative bacterial infections.

## Materials and Methods

### Key resources table

| Reagent type | Designation | Source or | Identifier | Additional |
| --- | --- | --- | --- | --- |

| **(species) or resource** |  | **reference** |  | **information** |
| --- | --- | --- | --- | --- |
| cell line (Homo-sapiens) | HEK293T | ATCC | CRL-3216,<br>RRID:<br>CVCL_0063 | Human embryonic kidney cells for transient transfection |
| Strain, strain background (*Pseudomonas aeruginosa*) | *Pseudomonas aeruginosa* (Schroeter) Migula | ATCC | Cat#: 27853 | Antibacterial activity test |
| Strain, strain background (*Pseudomonas aeruginosa*) | FADDI-PA022 | (33, 34) |  | Antibacterial activity test |
| Strain, strain background (*Pseudomonas aeruginosa*) | FADDI-PA025 | (23) |  | Antibacterial activity test |
| Strain, strain background (*Acinetobacter baumannii*) | *Acinetobacter baumannii* Bouvet and Grimont | ATCC | Cat#: 19606 | Antibacterial activity test |
| Strain, strain background (*Pseudomonas aeruginosa*) | FADDI-AB034 | (35) |  | Antibacterial activity test |
| Strain, strain background (*Acinetobacter baumannii*) | *Acinetobacter baumannii* Bouvet and Grimont | ATCC | Cat#: 17978 | Antibacterial activity test |
| Strain, strain background (*Klebsiella pneumoniae* subsp. *Pneumoniae*) | *Klebsiella pneumoniae* subsp. *pneumoniae* (Schroeter) Trevisan | ATCC | Cat#: 13883 | Antibacterial activity test |
| Strain, strain background<br>(*Klebsiella pneumoniae*) | FADDI-KP032 | (36) |  | Antibacterial activity test |
| Strain, strain background<br>(*Escherichia coli*) | FADDI-EC006 | (37) |  | Antibacterial activity test |
| Strain, strain background<br>(*Enterobacter cloacae*) | FADDI-EC001 | (37) |  | Antibacterial activity test |
| Strain, strain background<br>(*Enterobacter cloacae*) | FADDI-EC003 | (38) |  | Antibacterial activity test |
| Commercial assay or kit | QuickChange II site-directed mutagenesis kit | Agilent Technologies | Cat#: 200521 | Site-directed mutagenesis |
| Commercial assay or kit | BigDye™ Terminator v3.1 kit | Thermo Scientific | Cat#: 4337461 | DNA sequencing |
| Commercial assay or kit | Lipofectamine 2000 Reagent | Invitrogen | Cat#: 11668019 | Plasmid transfection |
| Commercial assay or kit | SlowFade™ Gold Antifade Mountant with DAPI | Invitrogen | Cat#: S36942 | Anti-fading for fluorescent probe and nuclear staining |
| Commercial assay or kit | Streptavidin-Agarose resin | Abcam | Cat#: ab270530 | Cell surface biotinylation assay |
| Commercial assay or kit | Sulfo-NHS-SS-Biotin | Campbell Science | Cat#: BR109 | Cell surface biotinylation assay |
| Commercial assay or kit | Immobilon Western Chemiluminesce nt HRP substrate | Merck Millipore | Cat#: WBKLS0500 | Western blot signal detection |
| Chemical compound, drug | Glycylsarcosine, [<sup>3</sup>H] | BioScientific Pty. Ltd | Cat#: MT 1545-250 | hPepT2 uptake assay substrate |
| Chemical compound, drug | Glycylsarcosine | Sapphire Biosciences | Cat#: BS-99594C | hPepT2 uptake assay substrate |
| Chemical compound, drug | Polymyxin B<sub>1</sub> | Sapphire Biosciences | Cat#: 14074 | hPepT2 uptake assay substrate |
| Chemical compound, drug | Polymyxin B sulfate | BetaPharm (Shanghai) | Cat#: C56H100N16O17S | hPepT2 uptake assay substrate and antibiotic used in animal study |
| Chemical compound, drug | MIPS-9541 | (13, 39, 40) |  | hPepT2 uptake assay substrate |
| Chemical compound, drug | FADDI-167 | (23, 41) |  | Polymyxin analogue |
| Chemical compound, drug | FADDI-170 | (23, 41) |  | Polymyxin analogue |
| Chemical compound, drug | FADDI-175 | (23, 41) |  | Polymyxin analogue |
| Chemical compound, drug | FADDI-793 | (23, 41) |  | Polymyxin analogue |
| Chemical compound, drug | FADDI-795 | (23, 41) |  | Polymyxin analogue |
| Antibody | Flag (DYKDDDDK) rabbit polyclonal antibody | Genesearch | Cat#: 2368, RRID: AB_2217020 | Western blot (1:1,000 dilution) |
| Antibody | β-actin mouse monoclonal antibody | Genesearch | Cat#: 4967, RRID: AB_330288 | Western blot (1:2,000 dilution) |
| Antibody | Goat anti-rabbit IgG conjugated with HRP 2<sup>nd</sup> | Sigma-Aldrich | Cat#: A0545, RRID: | Western blot (1:5,000 dilution) |
|  | antibody |  | AB_257896 |  |
| Antibody | Goat anti-mouse IgG conjugated with HRP 2<sup>nd</sup> antibody | Sigma-Aldrich | Cat#: A2554, RRID: AB_258008 | Western blot (1:10,000 dilution) |
| Other | Dulbecco's Modified Eagle Medium (DMEM) | Thermo Scientific | Cat#: 11995073 | Cell culture medium |
| Other | Glutamine | Thermo Scientific | Cat#: 25030081 | Cell culture supplement |
| Other | Fetal bovine serum (FBS) | Gibco | Cat#: 26140-079 | Cell culture supplement |
| Recombinant DNA reagent | hPepT2 | Australian Biosearch | Plasmid | PCMV6-XL5-hPepT2 |
| Recombinant DNA reagent | hPepT2-c-Flag | Australian Biosearch | Plasmid | pcDNA3.1+/C-(K)DYK-hPepT2-c-Flag |
| Software, algorithm | Clustal Omega | University College Dublin | RRID: SCR_001591 | Sequence Alignment |
| Software, algorithm | MODELLER | University of California, San Francisco | RRID: SCR_008395 | Model prediction |
| Software, algorithm | AlphaFold2 | Google DeepMind | RRID: SCR_025454 | Model prediction |
| Software, algorithm | Martini Maker module | CHARMM-GUI | RRID:SCR_025037 | Model construction |
| Software, algorithm | PyMOL | Schrodinger | Version 3.1.3.1, RRID: SCR_000305 | Molecular visualizations, structural figures, and trajectory animations |
| Software,<br>algorithm | ChimeraX | University of<br>California,<br>San<br>Francisco | Version 1.1,<br>RRID:<br>SCR_015872 | Molecular<br>visualizations,<br>structural figures,<br>and trajectory<br>animations |
| Software,<br>algorithm | AutoDock Vina | The Scripps<br>Research<br>Institute | RRID:<br>SCR_011958 | Molecular<br>docking |
| Software,<br>algorithm | CHARMM-GUI | Lehigh<br>University | RRID:<br>SCR_025037 | System<br>preparation |
| Software,<br>algorithm | GROMACS | University of<br>Groningen | Version 2021.2,<br>RRID:<br>SCR_014565 | Molecular<br>dynamic<br>simulations |
| Software,<br>algorithm | GraphPad Prism | GraphPad<br>Software | Version 10.0,<br>RRID:<br>SCR_002798 | Statistical<br>analysis |
| Software,<br>algorithm | ImageJ | National<br>Institutes of<br>Health | RRID:<br>SCR_003070 | Image processing<br>and<br>quantification |

### Reagents and chemicals

[^3^H] Gly-Sar (2 Ci/µmol) was obtained from BioScientific Pty. Ltd. (Gymea, NSW, Australia). Dulbecco’s Modified Eagle Medium (DMEM), fetal bovine serum (FBS), and other culture supplements were purchased from Thermo Scientific (Lidcombe, NSW, Australia). Gly-Sar and polymyxin B1 for molecular studies were purchased from Sapphire Biosciences (Redfern, NSW, Australia). Polymyxin B sulfate for animal studies was purchased from BetaPharm (Shanghai, China). MIPS-9541 was synthesised as previously reported (13, 39, 40). Plasmids containing the full-length hPepT2 and hPepT2-c-Flag were obtained from Australian Biosearch (Balcatta, WA, Australia). All other reagents and chemicals were purchased from Sigma-Aldrich (Castle Hill, NSW, Australia).

### Structural modelling of hPepT2 and polymyxins

The inward-open conformation of hPepT2 was predicted using AlphaFold2 via the ColabFold implementation with default parameters, employing the MMseqs2 multiple sequence alignment pipeline. To obtain the physiologically relevant outward-open conformation of hPepT2 for substrate binding, homology modeling was performed using MODELLER with the cryo-EM structure of rabbit PepT2 in the outward-open state (PDB: 7NQK) as the template. The sequence alignment between hPepT2 and rabbit PepT2 was conducted using Clustal Omega. One hundred models were generated, and the final model was selected based on the lowest Discrete Optimized Protein Energy score and verified by Ramachandran plot analysis (with >95% of residues in favored regions). The structure of polymyxin B1 was constructed and energy-minimized using the CHARMM36 force field. Computational alanine scanning of polymyxin B1 was performed by individually replacing each Dab (2,4-diaminobutyric acid) residue at positions 1, 3, 5, 8, and 9 with alanine using the mutagenesis wizard in PyMOL.

### Coarse-grained molecular dynamics (MD) simulations

Polymyxin B1 was employed as the prototype (27) and the coarse-grained (CG) model system of hPepT2 was constructed using the Martini Maker module in CHARMM-GUI (28). The membrane lipid composition was 50% phosphatidylcholine, 25% phosphatidylethanolamine, and 25% phosphatidylserine (29). In each simulation system, 10 coarse-grained polymyxin molecules were randomly introduced to enhance the probability of binding to hPepT2. These were solvated using CG water particles and neutralised with 0.1 M sodium chloride in a 15 × 15 × 14 nm^3^ simulation box. All CG simulations were conducted under periodic boundary conditions using the GROMACS program (version 2021.2) and the Martini 2.2 force field (30, 31). To eliminate steric interference, independent steepest energy minimization was performed for every system to achieve the maximum force below 1,000 kJ mol^-1^ nm^-2^. Subsequently, a six-step equilibration cycle was carried out by gradually turning off the position restraints on the lipid and polymyxin molecules. Finally, 3 µs production simulations were performed for each of two independent replicates (including independent energy minimization) were conducted for each system in the NPT (referring to the number of particles (N), system pressure (P), and temperature (T)) ensemble. Two replicates per condition were performed at this exploratory stage, as the CG-MD simulations were designed to identify the binding pathway of polymyxin B to hPepT2; the final binding poses from both replicates were highly consistent. Quantitative interaction analysis was subsequently performed using all-atom MD simulations with four replicates. The temperature and pressure were coupled to 310 K using the velocity rescaling method (time constant of 1 ps) and 1 bar using the semi-isotropic barostat and Parrinello-Rahman algorithm (time constant of 5 ps), respectively (32). Molecular visualizations, structural figures, and trajectory animations were prepared using PyMOL (Schrodinger) and ChimeraX. An animation of the CG-MD binding trajectory was provided as Video 1.

### All-atom MD simulations

The previous study of PepT_Xc_, a homolog of hPepT2, revealed that the protonation of its His67 promotes the outward-opening conformation (33); therefore, the corresponding residue His87 of hPepT2 was set to be protonated during all-atom simulations to bias the system towards an outward opening state. In the two CG-MD replicates, polymyxin B eventually bound to the lateral opening gate of hPepT2. To examine the detailed interaction between polymyxin and hPepT2, a single polymyxin B1 molecule was docked into the predicted binding pocket of hPepT2 using AutoDock Vina (34). Furthermore, to enhance the sampling of the binding dynamics, four different poses of polymyxin B1 were employed to build the initial hPepT2-polymyxin complex. Force field parameters for polymyxin B1 were generated using CGenFF. The membrane protein system was then constructed using the membrane builder module in CHARMM-GUI (35). The membrane lipid composition was 50% phosphatidylcholine, 25% phosphatidylethanolamine, and 25% phosphatidylserine, consistent with the CG-MD simulations. Each system was solvated with TIP3P water molecules in a rectangular box of 11 × 11 × 14 nm^3^ and neutralized with 0.15 M NaCl, yielding a total of approximately 158,000 atoms.

GROMACS 2021.2 was used to perform all MD simulations with the CHARMM36 all-atom force field (36). A time step of 2 fs was employed to integrate the Newtonian equations of motion, and all bonds involving hydrogen atoms were constrained. Long-range electrostatic interactions were treated using the Particle Mesh Ewald (PME) method with a switching function applied between 10 and 12 Å (37). Energy minimization was performed using the steepest descent algorithm. Subsequently, the system was equilibrated through a series of MD simulation steps in the NVT (referring to the number of particles (N), system volume (V), and temperature (T)) and NPT ensembles (310 K, 1 bar), during which positional restraints on membrane and protein atoms were gradually reduced. The root mean square deviations of the hPepT2-polymyxin B1 complex were calculated to assess simulation equilibration (Fig. 9). No restraints were applied in the final stage of equilibration and throughout the production run. The PME method was employed to treat long-range electrostatic interactions with a short-range cut-off of 1.2 nm, while the shifted Lennard-Jones potential algorithm was used to calculate van der Waals interactions with a general cut-off of 1.2 nm and a shifting cut-off of 1.0 nm (38, 39). The trajectory during production simulations was recorded every 10 ps. Production simulations were conducted for 500 ns for outward-open conformation sampling and 100 ns for polymyxin-hPepT2 interaction analysis.

**Figure 9.**
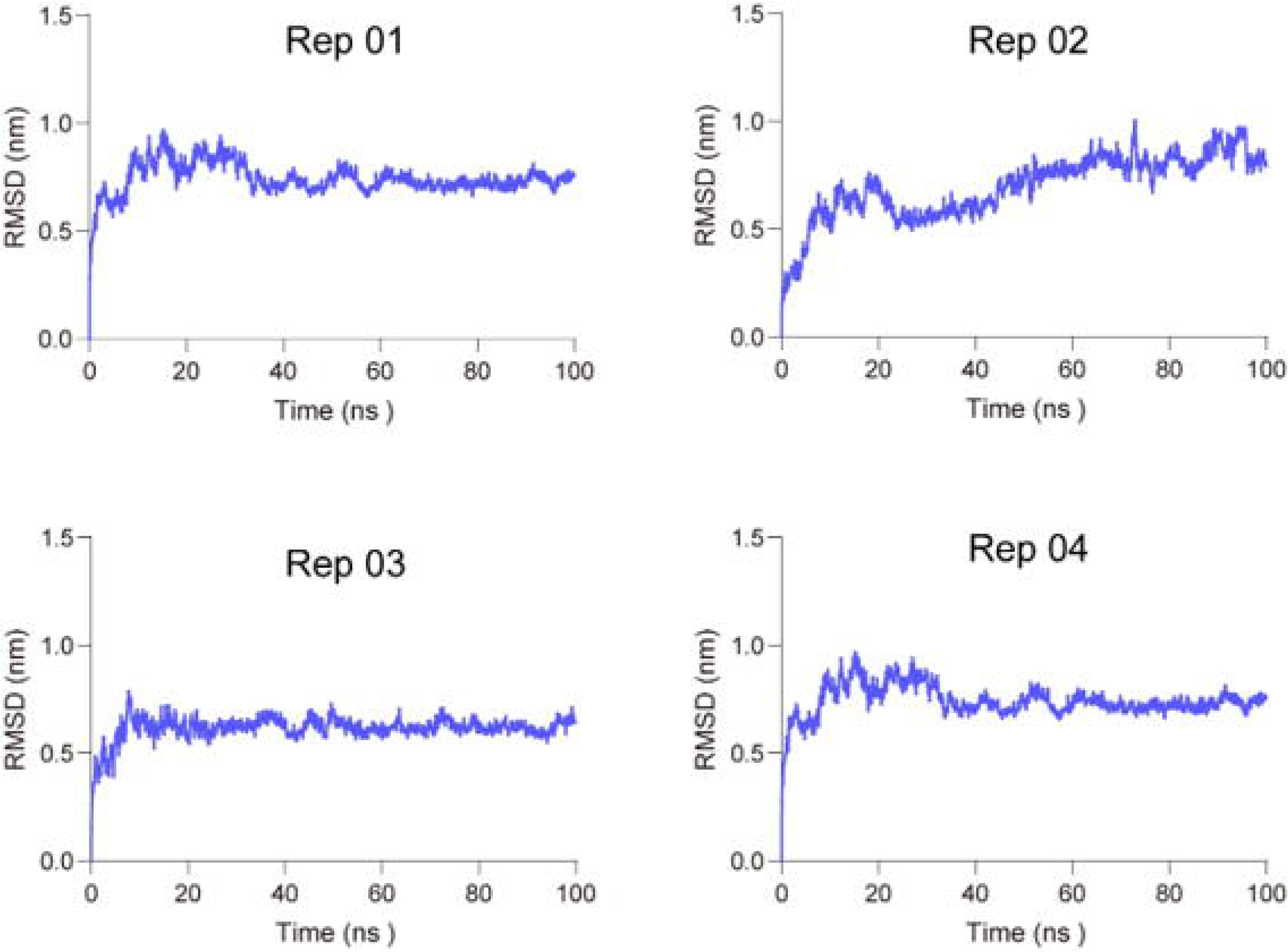
Time evolution of the root mean square deviations of polymyxin B. The results are calculated based on the four independent all-atom molecular dynamics simulation replicates.

### Site-directed mutagenesis and plasmid DNA sequencing

The hPepT2 mutants were generated using a QuickChange II site-directed mutagenesis kit (Agilent Technologies, Mulgrave, Victoria, Australia) with the primers listed in Table 4. All mutant construct sequences were confirmed by DNA sequencing (Ramaciotti Centre for Gene Function Analysis, Randwick, NSW, Australia) with the BigDye^TM^ Terminator v3.1 kit (Thermo Scientific, Lidcombe, NSW, Australia) (11, 27, 42–44).

### Transfection of hPepT2 and its mutants into HEK293 cells

HEK293T cells were cultured in DMEM supplemented with 10% FBS at 37°C with 5% CO_2_. HEK293T cell line was authenticated by the Australian Genome Research Facility (Urrbrae, SA, Australia) and checked for mycoplasma every 3 months. Cells were transfected with hPepT2 or its mutant plasmid DNA using Lipofectamine 2000 Reagent according to the manufacturer’s instructions (Invitrogen, Mount Waverley, VIC, Australia) (9, 13, 28, 45, 46). At 24 h after transfection, the cellular uptake of [^3^H] Gly-Sar or MIPS-9541 was assessed.

### Transport uptake assay

To evaluate the uptake of [^3^H] Gly-Sar via hPepT2 or its mutant transporters, we measured the accumulation of [^3^H] Gly-Sar in hPepT2-overexpressing HEK293 cells. The uptake was conducted at 37°C in phosphate-buffered saline (PBS, pH 5.0) containing 5 mM glucose. The specific substrate concentration was 2.5 μM [^3^H] Gly-Sar, and the duration was 8 min according to our previous studies (9, 13). The uptake assay was terminated by three rapid washes with ice-cold PBS. Samples were lysed in 0.2 M NaOH, neutralized in 0.2 M HCL and then processed for liquid scintillation counting. The uptake counts for hPepT2 or its mutant expressing cells were all subtracted by those of the control.

To evaluate the uptake of polymyxin B via hPepT2 or its mutant transporters, we measured the accumulation of a validated fluorescent polymyxin probe MIPS-9541 and polymyxin analogues (i.e. FADDI-167, -170, -175, -793, and -795, Fig. 10) in hPepT2-overexpressing HEK293 cells (13). The uptake was performed in PBS (pH 5.0) supplemented with 5 mM glucose for 10 min at 37°C. Intracellular fluorescence accumulation was quantified with excitation/emission wavelengths of 350 nm/518 nm using a Tecan Safire II microplate reader (Thermo Scientific, Lidcombe, NSW, Australia). Background counts of the vector-transfected cells were subtracted from all uptake measurements. Kinetic analysis was conducted by measuring the uptake of transporter-expressing cells with a range of concentrations of MIPS-9541 (0 - 50 µM) over 10 min. Apparent K_m_ and V_max_ values were determined using GraphPad Prism 10.0 (13, 47).

**Figure 10.**
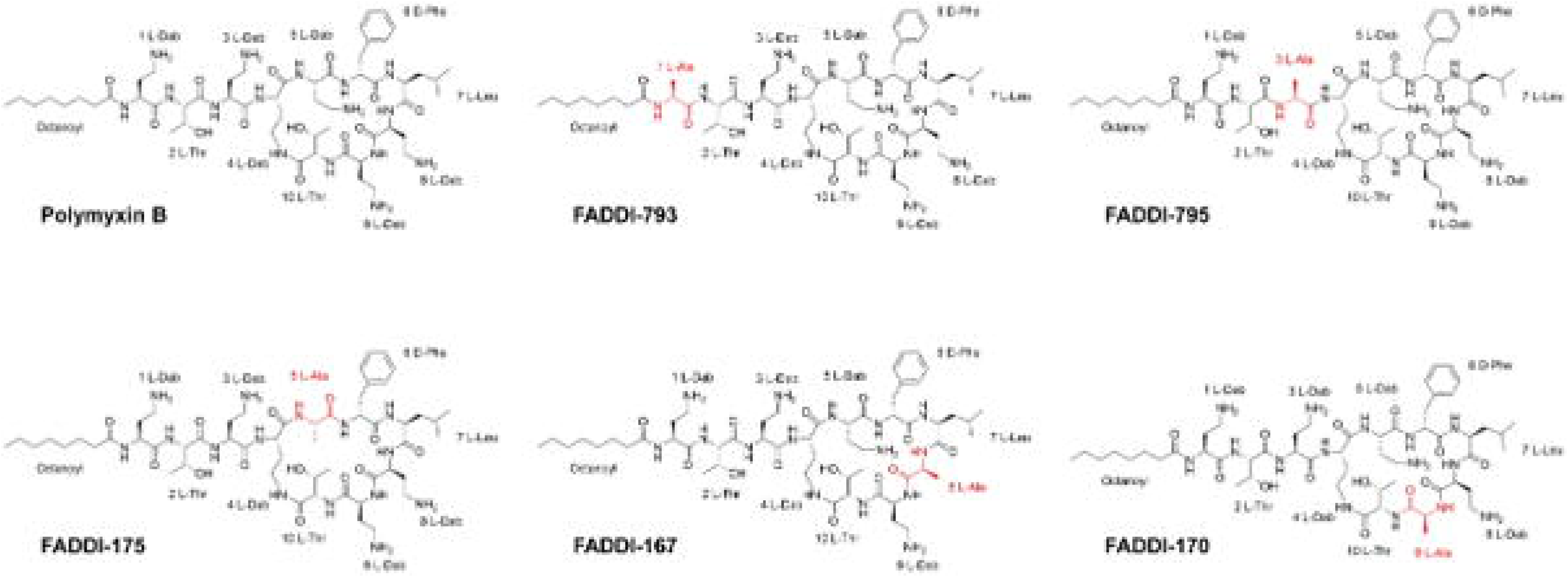
Chemical structures of polymyxin B and its analogues. The modified structural moieties are labelled in red.

### Fluorescence imaging of MIPS-9541

HEK293 cells were seeded on chamber slides and transfected with hPepT2 or its mutant plasmids. After 24 h, the culture medium was removed and cells were incubated with freshly prepared MIPS-9541 solution in PBS (10 µM, pH 5.0) for 10 min at 37°C. Cells were then washed with ice-cold PBS three times and mounted in SlowFade Gold Antifade Mountant reagent supplemented with 4’,6-diamidino-2-phenylindole (DAPI, Thermo Scientific, Lidcombe, NSW, Australia). Samples were imaged with a Leica Thunder 3D Imager (Leica Microsystems, North Ryde, NSW, Australia).

### Cell surface biotinylation

We employed sulfo-NHS-SS-Biotin (Campbell Science, Rockford, IL, USA) to label the plasma membrane proteins in HEK293 cells overexpressing hPepT2 or its mutants (48). Cell culture plates were pre-chilled on ice. Culture medium was aspirated, and cells were washed twice with cold PBS (pH 8.0). Freshly prepared 1 mg/mL sulfo-NHS-SS-Biotin in PBS (pH 8.0) was then added to each well and incubated for 30 min. Cells were washed twice with 100 mM glycine in PBS (pH 8.0) and then three times with PBS (pH 7.4). Cells were lysed in the lysis buffer (Tris 10 mM, NaCl 150 mM, EDTA 1 mM, SDS 0.1%, and Triton X-100 1% with 1:1,000 dilution of a protease inhibitor cocktail) (27). Cell lysate was subjected to centrifugation at 14,000 *g* and 4°C for 10 min, and the supernatant was collected. Pre-washed Streptavidin-Agarose beads were supplied to the samples and incubated with rotation at 4°C for 1 h. After three rounds of centrifugation and washing with PBS (pH 7.4), beads were resuspended in PBS with 1x Laemmli buffer containing 2.5% β-mercaptoethanol and incubated at 55°C for 30 min. The remaining supernatant was collected, denatured, and subjected to electrophoresis.

### Total cell lysis, electrophoresis and immunoblotting

Upon washing, cells were collected and lysed in the lysis buffer. After incubating on ice for 10 min, cell lysate was centrifuged at 14,000 *g* and 4°C for 10 min. The supernatant was collected as total cell lysate and denatured at 55°C in 1x Laemmli buffer containing 2.5% β-mercaptoethanol for 30 min.

Protein lysate samples were loaded onto 10% SDS-PAGE gels followed by electrophoresis as described before (49), and the protein samples on the gels were transferred to polyvinylidene difluoride (PVDF) membranes. Following blocking with 5% non-fat milk in PBS-Tween (Na_2_HPO_4_ 80 mM, KH_2_PO_4_ 20 mM, NaCl 100 mM, and 0.05% Tween 20, pH 7.5) and multiple washings with PBS-Tween, the immunoblots were incubated at 4°C overnight with anti-flag (DYKDDDDK) antibody (1:1,000; Catalogue Number 2368, Genesearch, Arundel, QLD, Australia) or anti-β-actin antibody (1:2,000, Catalogue Number 4967, Genesearch). After washing with PBS-Tween three times, the immunoblots were incubated at room temperature for 1 h with goat anti-rabbit IgG conjugated with HRP (1:5,000; Catalogue Number A0545, Sigma-Aldrich) or goat anti-mouse IgG conjugated with HRP (1:10,000; Catalogue Number A2554, Sigma-Aldrich). The blots were washed, incubated with Immobilon Western Chemiluminescent HRP substrate (Merck Millipore, Kilsyth, VIC, Australia), and imaged with an ImageQuant LAS 500 (GE Healthcare, Avantor, PA, USA). The densitometry analysis of immunoblots was processed with ImageJ (NIH, Bethesda, MD, USA).

### Chemical synthesis of polymyxin analogues

Five lipopeptides (FADDI-167, FADDI-170, FADDI-175, FADDI-793, and FADDI-795) were synthesised and purified in house as described before (23, 41). Briefly, the lipopeptides were prepared using a Protein Technologies Prelude automated peptide synthesizer according to the standard protocol for Fmoc solid-phase peptide chemistry.

### Measurements of minimum inhibitory concentrations

MICs were measured using the broth micro-dilution method with clinical isolates and American Type Culture Collection (ATCC) strains (37). Each bacterial strain (∼10^6^ colony-forming units [CFU] per mL) was suspended in 100 μL of freshly prepared cation-adjusted Mueller-Hinton broth (CaMHB) in 96-well plates. Lipopeptides were diluted into different concentrations in CaMHB and added to the plates for subsequent incubation at 37°C for 18-24 h. MICs were determined as the lowest concentration at which the visible growth of bacteria was inhibited.

### Nephrotoxicity in mice

The ARRIVE guideline was followed for *in vivo* experiment reporting. The mouse nephrotoxicity study was approved by the Monash Animal Ethics Committee. Mice (Swiss, female, 6-8 weeks old) were maintained in micro-isolators in a temperature-controlled PC2 animal laboratory with an ambient humidity of 50-70% and a 12 h/12 h dark/light cycle. Mice were randomly divided into groups. Lipopeptide solutions were subcutaneously administered to mice with a dose of 8 mg/kg every 2 h for 6 doses in one day (a total dose of 48 mg/kg/day, n=3 each group). At 24 h after the last injection, mice were sacrificed, the kidneys were collected from mice immediately, and fixed in 10% formalin buffer (pH 7.4, Sigma-Aldrich, Australia). Histological examination of the kidney samples was conducted at the Australian Phenomics Network-Histopathology and Organ Pathology (University of Melbourne, Parkville, VIC, Australia). An experienced pathologist blinded to treatment groups assessed renal histology and graded the severity of nephrotoxicity using a semi-quantitative scoring system (scores 1-10) as previously described (23). The percentage of kidney affected was also scaled in the range of 0-6 (0: <1%; 1: 1 to <5%; 2: 5 to <10%; 3: 10 to <20%; 4: 20 to <30%; 5: 30 to ≤40% and 6: >40%). A semi-quantitative score (SQS) was calculated using the grade score and kidney damage percentage to quantify the overall kidney damage (23). Noteworthy, in this proof-of-concept study, we employed acute kidney damage analysis in a small number of mice to rapidly screen these analogues for nephrotoxicity. Three biological replicates were used because our mouse nephrotoxicity model is robust as demonstrated in our drug discovery program (23). Moreover, minimising animal use is a fundamental principle of the 3Rs (Replacement, Reduction, and Refinement).

### Statistical analyses

Unpaired *t*-test was employed to detect the variance between the protein expression of hPepT2 and its mutants. Welch’s *t*-test was used to assess the differences between the transport function or kinetic parameters of hPepT2 and its mutants. All data are presented as mean ± standard deviation (SD), and a *p* value <0.05 was considered statistically significant. All experiments were repeated three times with triplicates in each experiment.

## Acknowledgements

Not applicable.

## Funding

We acknowledge the funding support of the e-Asia Joint Research Program (2018589). XJ is supported by the National Natural Science Foundation of China (32401034, 32301041, and 32571437), Basic Research Program of Jiangsu (BK20240425), and the Shandong Excellent Young Scientists (Overseas) Fund Program (2023HWYQ-044). VT is supported by the Program Management Unit B (Brain Power, Manpower), Ministry of Higher Education, Science, Research and Innovation, Thailand. JL is an Australian National Health and Medical Research Council (NHMRC) Investigator Fellow (APP2025937).

## Data availability

All strains involved in this study are available upon request. All data analysed in the current study are provided in the main text and supporting files.

## Declarations

### Ethics approval and consent to participate

This study was approved by the Monash Animal Ethics Committee before commencement (AEC37419).

### Consent for publication

Not applicable.

### Competing interests

JL, TV, and KDR are listed as inventors on the patent application WO2015149131 ‘Polymyxin Derivatives as Antimicrobial Compounds’ which covers new-generation polymyxins and has been licensed to Qpex Biopharma and Brii Biosciences. JL received grants, speaking honoraria, and consulting fees from Northern Antibiotics, Avexa, Genentech, Healcare, CTTQ, Aosaikang, Jiayou Medicine, MedCom, Fansheng Biotech, DanDi BioScience, Qpex Biopharma, and Xellia Pharmaceuticals. JL is CEO of Sliabx Pharmaceuticals and Cinfinno Biotech and holds equities in both companies. All other authors declare no conflict of interest.

## Abbreviations

ATCC: American Type Culture Collection
CaMHB: cation-adjusted Mueller-Hinton broth
CG: coarse-grained
Coul: electrostatic interaction
Cryo-EM: Cryo-electron microscopy
DAPI: 4’,6-diamidino-2-phenylindole
DMEM: Dulbecco’s Modified Eagle Medium
ECD: extracellular domain
FBS: fetal bovine serum
Gly-Sar: glycosarcosine
hPepT2: human oligopeptide transporter 2
MD: molecular dynamics
MDR: multidrug-resistant
MIC: minimum inhibitory concentration
NPT: number of particles, system pressure and temperature
NVT: number of particles, system volume and temperature
PBS: phosphate-buffered saline
PME: Particle Mesh Ewald
PVDF: polyvinylidene difluoride
SD: standard deviation
SIR: structure-interaction relationship
SLCs: solute carrier transporters
SQS: semi-quantitative score
TM: transmembrane domain
VDW: hydrophobic interaction

## Notes

### Summary of Updates

We have made some additional minor changes according to the comments from the editorial office: (1) integrating supplementary figures into the main text (2) categorizing the video file as a rich media file, (3) providing the required information in the Key Resources Table, (4) removing the author contributions, and (5) listing the gene name appropriately.

## References

1. Ardal C, Balasegaram M, Laxminarayan R, McAdams D, Outterson K, Rex JH, et al. Antibiotic development - economic, regulatory and societal challenges. Nat Rev Microbiol. 2020;18(5):267–74.

2. Collaborators GBDAR. Global burden of bacterial antimicrobial resistance 1990-2021: a systematic analysis with forecasts to 2050. Lancet. 2024;404(10459):1199–226.

3. Morehead MS, Scarbrough C. Emergence of global antibiotic resistance. Prim Care. 2018;45(3):467–84.

4. Baindara P, Kumari S, Dinata R, Mandal SM. Antimicrobial peptides: evolving soldiers in the battle against drug-resistant superbugs. Mol Biol Rep. 2025;52(1):432.

5. Nang SC, Azad MAK, Velkov T, Zhou QT, Li J. Rescuing the last-line polymyxins: achievements and challenges. Pharmacol Rev. 2021;73(2):679–728.

6. Azad MAK, Nation RL, Velkov T, Li J. Mechanisms of polymyxin-induced nephrotoxicity. Adv Exp Med Biol. 2019;1145:305–19.

7. Wang JL, Xiang BX, Song XL, Que RM, Zuo XC, Xie YL. Prevalence of polymyxin-induced nephrotoxicity and its predictors in critically ill adult patients: A meta-analysis. World J Clin Cases. 2022;10(31):11466–85.

8. Slingerland CJ, Martin NI. Recent advances in the development of polymyxin antibiotics: 2010-2023. ACS Infect Dis. 2024;10(4):1056–79.

9. Lu X, Chan T, Zhu L, Bao X, Velkov T, Zhou QT, et al. The inhibitory effects of eighteen front-line antibiotics on the substrate uptake mediated by human Organic anion/cation transporters, Organic anion transporting polypeptides and Oligopeptide transporters in in vitro models. Eur J Pharm Sci. 2018;115:132–43.

10. Zhou F, Zhu L, Wang K, Murray M. Recent advance in the pharmacogenomics of human Solute Carrier Transporters (SLCs) in drug disposition. Adv Drug Deliv Rev. 2017;116:21–36.

11. Chan T, Lu X, Shams T, Zhu L, Murray M, Zhou F. The role of N-glycosylation in maintaining the transporter activity and expression of human oligopeptide transporter 1. Mol Pharm. 2016;13(10):3449–56.

12. Luo Y, Gao J, Jiang X, Zhu L, Zhou QT, Murray M, et al. Molecular insights to the structure-interaction relationships of human proton-coupled oligopeptide transporters (PepTs). Pharmaceutics. 2023;15(10):2517.

13. Lu X, Chan T, Xu C, Zhu L, Zhou QT, Roberts KD, et al. Human oligopeptide transporter 2 (PEPT2) mediates cellular uptake of polymyxins. J Antimicrob Chemother. 2016;71(2):403–12.

14. Ocheltree SM, Shen H, Hu Y, Xiang J, Keep RF, Smith DE. Role of PEPT2 in the choroid plexus uptake of glycylsarcosine and 5-aminolevulinic acid: studies in wild-type and null mice. Pharm Res. 2004;21(9):1680–5.

15. Smith DE, Clemencon B, Hediger MA. Proton-coupled oligopeptide transporter family SLC15: physiological, pharmacological and pathological implications. Mol Aspects Med. 2013;34(2-3):323–36.

16. Zhao D, Lu K. Substrates of the human oligopeptide transporter hPEPT2. Biosci Trends. 2015;9(4):207–13.

17. Xiang J, Chiang PP, Hu Y, Smith DE, Keep RF. Role of PEPT2 in glycylsarcosine transport in astrocyte and glioma cultures. Neurosci Lett. 2006;396(3):225–9.

18. Alghamdi OA, King N, Jones GL, Moens PDJ. A new use of beta-Ala-Lys (AMCA) as a transport reporter for PEPT1 and PEPT2 in renal brush border membrane vesicles from the outer cortex and outer medulla. Biochim Biophys Acta Biomembr. 2018;1860(5):960–4.

19. Shen H, Keep RF, Hu Y, Smith DE. PEPT2 (Slc15a2)-mediated unidirectional transport of cefadroxil from cerebrospinal fluid into choroid plexus. J Pharmacol Exp Ther. 2005;315(3):1101–8.

20. Ganapathy ME, Huang W, Wang H, Ganapathy V, Leibach FH. Valacyclovir: a substrate for the intestinal and renal peptide transporters PEPT1 and PEPT2. Biochem Biophys Res Commun. 1998;246(2):470–5.

21. Newstead S, Parker J, Deme J, Lichtinger S, Kuteyi G, Biggin P, et al. Structural basis for antibiotic transport and inhibition in PepT2, the mammalian proton-coupled peptide transporter. Res Sq. 2024:rs.3.rs-4435259.

22. Parker JL, Deme JC, Wu Z, Kuteyi G, Huo J, Owens RJ, et al. Cryo-EM structure of PepT2 reveals structural basis for proton-coupled peptide and prodrug transport in mammals. Sci Adv. 2021;7(35):eabh3355.

23. Roberts KD, Zhu Y, Azad MAK, Han ML, Wang J, Wang L, et al. A synthetic lipopeptide targeting top-priority multidrug-resistant Gram-negative pathogens. Nat Commun. 2022;13(1):1625.

24. Li J, Nation RL, Turnidge JD, Milne RW, Coulthard K, Rayner CR, et al. Colistin: the re-emerging antibiotic for multidrug-resistant Gram-negative bacterial infections. Lancet Infect Dis. 2006;6(9):589–601.

25. Abdelraouf K, Chang KT, Yin T, Hu M, Tam VH. Uptake of polymyxin B into renal cells. Antimicrob Agents Chemother. 2014;58(7):4200–2.

26. Jiang X, Zhang S, Azad MAK, Roberts KD, Wan L, Gong B, et al. Structure-interaction relationship of polymyxins with the membrane of human kidney proximal tubular cells. ACS Infect Dis. 2020;6(8):2110–9.

27. Chan T, Zheng J, Zhu L, Grewal T, Murray M, Zhou F. Putative transmembrane domain 6 of the human organic anion transporting polypeptide 1A2 (OATP1A2) influences transporter substrate binding, protein trafficking, and quality control. Mol Pharm. 2015;12(1):111–9.

28. Chan T, Cheung FS, Zheng J, Lu X, Zhu L, Grewal T, et al. Casein Kinase 2 Is a Novel Regulator of the Human Organic Anion Transporting Polypeptide 1A2 (OATP1A2) Trafficking. Mol Pharm. 2016;13(1):144–54.

29. Shen J, Hu M, Fan X, Ren Z, Portioli C, Yan X, et al. Extracellular domain of PepT1 interacts with TM1 to facilitate substrate transport. Structure. 2022;30(7):1035–41 e3.

30. Hong M, Li S, Zhou F, Thomas PE, You G. Putative transmembrane domain 12 of the human organic anion transporter hOAT1 determines transporter stability and maturation efficiency. J Pharmacol Exp Ther. 2010;332(2):650–8.

31. Hong M, Zhou F, Lee K, You G. The putative transmembrane segment 7 of human organic anion transporter hOAT1 dictates transporter substrate binding and stability. J Pharmacol Exp Ther. 2007;320(3):1209–15.

32. Xu L, Haworth IS, Kulkarni AA, Bolger MB, Davies DL. Mutagenesis and cysteine scanning of transmembrane domain 10 of the human dipeptide transporter. Pharm Res. 2009;26(10):2358–66.

33. Lin YW, Zhou Q, Onufrak NJ, Wirth V, Chen K, Wang J, et al. Aerosolized polymyxin B for treatment of respiratory tract infections: determination of pharmacokinetic-pharmacodynamic indices for aerosolized polymyxin B against pseudomonas aeruginosa in a mouse lung infection model. Antimicrob Agents Chemother. 2017;61(8):e00211–17.

34. Lin YW, Zhou QT, Cheah SE, Zhao J, Chen K, Wang J, et al. Pharmacokinetics/pharmacodynamics of pulmonary delivery of colistin against pseudomonas aeruginosa in a mouse lung infection model. Antimicrob Agents Chemother. 2017;61(3):e02025–16.

35. Kumar G. Natural peptides and their synthetic congeners acting against Acinetobacter baumannii through the membrane and cell wall: latest progress. RSC Med Chem. 2025;16(2):561–604.

36. Deris ZZ, Yu HH, Davis K, Soon RL, Jacob J, Ku CK, et al. The combination of colistin and doripenem is synergistic against Klebsiella pneumoniae at multiple inocula and suppresses colistin resistance in an in vitro pharmacokinetic/pharmacodynamic model. Antimicrob Agents Chemother. 2012;56(10):5103–12.

37. Roberts KD, Azad MA, Wang J, Horne AS, Thompson PE, Nation RL, et al. Antimicrobial activity and toxicity of the major lipopeptide components of polymyxin B and colistin: last-line antibiotics against multidrug-resistant Gram-negative bacteria. ACS Infect Dis. 2015;1(11):568–75.

38. Thombare VJ, Swarbrick JD, Azad MAK, Zhu Y, Lu J, Yu HY, et al. Exploring structure-activity relationships and modes of action of laterocidine. ACS Cent Sci. 2024;10(9):1703–17.

39. Deris ZZ, Swarbrick JD, Roberts KD, Azad MA, Akter J, Horne AS, et al. Probing the penetration of antimicrobial polymyxin lipopeptides into gram-negative bacteria. Bioconjug Chem. 2014;25(4):750–60.

40. Yun B, Azad MA, Nowell CJ, Nation RL, Thompson PE, Roberts KD, et al. Cellular uptake and localization of polymyxins in renal tubular cells using rationally designed fluorescent probes. Antimicrob Agents Chemother. 2015;59(12):7489–96.

41. Jiang X, Patil NA, Azad MAK, Wickremasinghe H, Yu H, Zhao J, et al. A novel chemical biology and computational approach to expedite the discovery of new-generation polymyxins against life-threatening Acinetobacter baumannii. Chem Sci. 2021;12(36):12211–20.

42. Zheng J, Chan T, Cheung FS, Zhu L, Murray M, Zhou F. PDZK1 and NHERF1 regulate the function of human organic anion transporting polypeptide 1A2 (OATP1A2) by modulating its subcellular trafficking and stability. PLoS One. 2014;9(4):e94712.

43. Zhou F, Zheng J, Zhu L, Jodal A, Cui PH, Wong M, et al. Functional analysis of novel polymorphisms in the human SLCO1A2 gene that encodes the transporter OATP1A2. AAPS J. 2013;15(4):1099–108.

44. Zhou F, Zhu L, Cui PH, Church WB, Murray M. Functional characterization of nonsynonymous single nucleotide polymorphisms in the human organic anion transporter 4 (hOAT4). Br J Pharmacol. 2010;159(2):419–27.

45. Ali Y, Shams T, Cheng Z, Li Y, Chun CS, Shu W, et al. Impaired transport activity of human Organic anion transporters (OATs) and Organic anion transporting polypeptides (OATPs) by Wnt inhibitors. J Pharm Sci. 2021;110(2):914–24.

46. Li Z, Cheung FS, Zheng J, Chan T, Zhu L, Zhou F. Interaction of the bioactive flavonol, icariin, with the essential human solute carrier transporters. J Biochem Mol Toxicol. 2014;28(2):91–7.

47. Chan T, Zhu L, Madigan MC, Wang K, Shen W, Gillies MC, et al. Human organic anion transporting polypeptide 1A2 (OATP1A2) mediates cellular uptake of all-trans-retinol in human retinal pigmented epithelial cells. Br J Pharmacol. 2015;172(9):2343–53.

48. Zhou F, Hong M, You G. Regulation of human organic anion transporter 4 by progesterone and protein kinase C in human placental BeWo cells. Am J Physiol Endocrinol Metab. 2007;293(1):E57–61.

49. Zhou F, Xu W, Tanaka K, You G. Comparison of the interaction of human organic anion transporter hOAT4 with PDZ proteins between kidney cells and placental cells. Pharm Res. 2008;25(2):475–80.

